# Biased removal and loading of centromeric histone H3 during reproduction underlies uniparental genome elimination

**DOI:** 10.1101/2021.02.24.432754

**Authors:** Mohan P.A. Marimuthu, Ravi Maruthachalam, Ramesh Bondada, Sundaram Kuppu, Ek-Han Tan, Anne Britt, Simon S.W. Chan, Luca Comai

## Abstract

Uniparental genome elimination is a dramatic case of centromeric failure, resulting in the postzygotic loss of a parental chromosome set. Genome partitioning during cell division relies on centromere attachment to spindle fibers through kinetochores. Centromeres are epigenetically specified by CENP-A (CENH3), a conserved centromeric specific histone H3 variant. In Arabidopsis, CENH3 modification results in haploid inducers, whose genome is eliminated frequently when crossed to the wild type. To investigate the underlying mechanism, we dissected the timing and molecular features of genome elimination. In zygotes and early embryos from genome elimination crosses, CENH3 occupied only the centromeres contributed by the wild-type parent. Haploid inducer chromosomes had defective kinetochores and missegregated, often forming micronuclei. This uniparental loss of centromere identity is initiated by the removal of altered CENH3 at fertilization, while wild-type CENH3 persists and maintains strong centromeric identity. Weak centromeres were capable of rebuilding functional kinetochores, but often failed when in competition with normal ones. We induced a similar weak state by mitotic dilution of wild-type CENH3. Furthermore, weakness was suppressed by crosses of haploid inducers to other variants of haploid inducers, and enhanced by mutations in *VIM1*, a ubiquitin ligase known to modify CENH3 and centromeric DNA methylation.. The differential stability of altered CENH3 during reproduction has important genetic and evolutionary implications.

## Introduction

Centromeres are epigenetic loci that assemble kinetochores and mediate spindle-microtubule-based chromosome segregation during cell division (Fukagawa and Earnshaw, 2014). In a majority of eukaryotes, chromatin harboring the centromeric histone H3 (CENP-A, aka CENH3 or HTR12 in *A.thaliana*), rather than the underlying DNA sequence, defines the centromere (Black and Cleveland, 2011; Nasuda et al., 2005; Talbert et al., 2002; Voullaire et al., 1993). Incorporation of CENP-A into chromatin is the sufficient and principal epigenetic factor in specifying centromere location on a chromosome (Fachinetti et al., 2013; Mendiburo et al., 2011) with few exceptions (Akiyoshi and Gull, 2014; Drinnenberg et al., 2016). After each replication, pre-existing CENP-A nucleosomes on the centromeric chromatin mediate replenishment and perpetuation of the centromere (Stellfox et al., 2013). Failure of CENP-A incorporation, or its depletion, leads to chromosome segregation defects, genome instability and lethality (Buchwitz et al., 1999; Fachinetti et al., 2013; Howman et al., 2000; Lermontova et al., 2011a; Ly et al., 2017, 2019; Régnier et al., 2005; Sanei et al., 2011). CENP-A/CENH3 consists of a hypervariable, unstructured amino-terminal domain and of a more conserved, but rapidly evolving histone fold domain (HFD) (Henikoff et al., 2001), whose interactions with chaperones facilitate loading in centromeric chromatin (Barnhart et al., 2011; Black et al., 2007a, 2007b; Camahort et al., 2007; Chen et al., 2015; Fachinetti et al., 2013; Fujita et al., 2007; Lermontova et al., 2013; Shelby et al., 1997). In Arabidopsis, CENH3 is expressed in dividing and in reproductive cells (Aw et al., 2010; Heckmann et al., 2011) where it loads on centromeres during G2 phase (Lermontova et al., 2011b). In contrast to regular histones, CENP-A is highly stable once incorporated into the centromeres, displaying negligible turnover (Bodor et al., 2013; Lermontova et al., 2011b; Smoak et al., 2016). Loss of CENP-A or CENH3 has been observed in actively dividing tissues such as during male gamete formation in *C. elegans* (Gassmann et al., 2012) and in terminally differentiated cells (Lee et al., 2010; McGregor et al., 2014; Swartz et al., 2019), including the vegetative cell of pollen (Schoft et al., 2009). In wild-type *A. thaliana,* both N- and C-terminal GFP-CENH3 fusions localize to the centromeres (De Storme et al., 2016; Ravi et al., 2010). The C-terminal fusion is removed from centromeres of the egg and central cells and from male centromeres soon after fertilization (Ingouff et al., 2010), only to be reloaded before the first zygotic mitosis. Similarly, removal and reloading has been observed during male meiosis (Lermontova et al., 2011a; Ravi et al., 2011). However, it is not clear whether fusions represent faithfully the behavior of the native protein.

Uniparental <u>g</u>enome <u>e</u>limination (GE), manifested cytologically by chromosome missegregation, results in uniparental haploids when certain genotypes are crossed (Fujiwara et al., 1997; Gibeaux et al., 2018; Ishii et al., 2016). Haploids are valued in plant breeding because, upon genome doubling, they form homozygous individuals, bypassing a lengthy inbreeding process (Dunwell, 2010). The molecular mechanisms responsible for GE are unknown. Uniparental loss of CENH3 triggered by centromere inactivity and asynchronous cell cycle between parental genomes (Sanei et al., 2011), persistent sister-chromatid cohesion (Ishii et al., 2010, 2016), or differences in centromere size (Wang and Dawe, 2018), perhaps resembling meiotic drive (Akera et al., 2017; Chmátal et al., 2014; Henikoff et al., 2001) could be responsible. In crosses leading to GE, conflict in centromeric maintenance between parental chromosomes could result from genetic and epigenetic differences (Ishii et al., 2016). In Arabidopsis, isogenic parents that differed only in CENH3 produced progeny displayed GE (Ravi and Chan, 2010). From this and other studies, uniparental centromere malfunction emerges as a probable cause of parent-specific GE (Brown et al., 2011; Gibeaux et al., 2018; Ishii et al., 2016; Metcalfe et al., 2007; Raychaudhuri et al., 2012; Sanei et al., 2011; Wang and Dawe, 2018). Complementation of a null *cenh3* mutation with a translational fusion incorporating GFP, the histone H3.3 N-terminal tail and the CENH3 HFD (GFP-ts, Figures 1A and 1B), results in a **h**aploid **i**nducer (HI): when this GFP-ts plant is crossed to the wild type (WT), the GFP-ts chromosomes are eliminated at a high frequency, resulting in haploid progeny. The process is about ten-fold more efficient when the GFP-ts plant is used as a female rather than a male (Ravi and Chan, 2010). The outcome of this haploid induction cross (hereafter GE cross) is not uniform: it also produces normal diploid seeds, aneuploid seeds, and aborted seeds (Figure 1B). We assume that HI involves fertilization followed by GE because aneuploids display residual HI chromosomes that are often severely damaged (Tan et al., 2015). Therefore, zygotes must inherit a functioning chromosome set from the wild-type, but a centromeric incompetent set from the HI parent (Tan et al., 2015). Frequently, these chromosomes undergo complex rearrangements (Tan et al., 2015) comparable to chromoanagenesis phenomena described in animal cells (Ly et al., 2017; Stephens et al., 2011). A grossly altered CENH3 such as GFP-ts is not a requirement for GE since it can also result when complementing the arabidopsis *cenh3* null mutants with CENH3 from other plant species (Maheshwari et al., 2015, 2017), or when one or two HFD AA residues have been substituted (Karimi-Ashtiyani et al., 2015; Kuppu et al., 2015, 2020) (Figure 1A). Wheat and maize heterozygous, respectively, for a large restored frameshift mutation in the N-terminus, and a null *cenh3* mutation, are efficient HI (Lv et al., 2020; Wang et al., 2021). It is unclear, however, what shared feature of the different inducers, if any, is responsible for GE.

**Figure 1.**
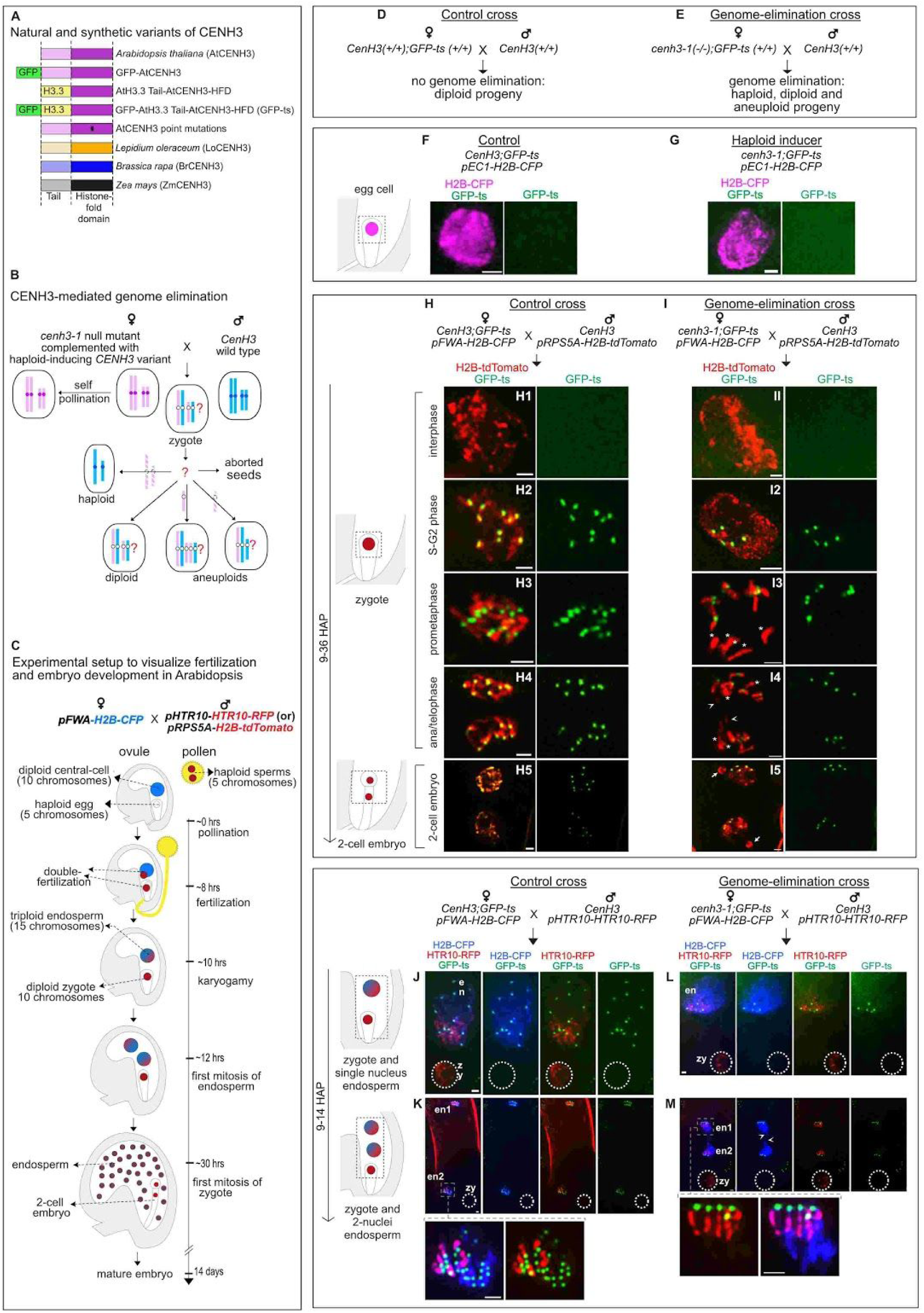
Uniparental localization of maternal GFP-ts during haploid induction. (A) Structure of wild-type AtCENH3 and haploid inducing CENH3 variants. All variants act as recessive alleles. (B) CENH3-mediated GE in Arabidopsis. “?”: unknown steps and mechanisms. (C) Experimental setup to image double-fertilization and early seed development in Arabidopsis. (D, E) Parental genotypes and outcome of control and genome-elimination crosses. The ovule schematic on the left indicates the region of interest in the ovule shown on the right (F-M). (F and G) Egg nucleus before fertilization. Progressive stages of zygotic development in control (H1-H5) and GE cross (I1-I5). “*” marks condensed chromosomes without centromeric GFP-ts (I3, I4); arrowheads: lagging chromosomes (I4); arrows: micronuclei (I5). Endosperm nuclei displaying male (red) and female (blue) chromatin in interphase (J, L) and metaphase (K) or anaphase chromatin (M). Arrowheads: female chromatin bridge (M). White dotted circles (J,K,L,M): zygote. en: endosperm; zy: zygote; HAP: hours after pollination. Scale bars, 1 µm.

Interestingly, HIs are compatible when selfed, yielding diploid progeny (Figure 1B), and exhibit incompatibility only when outcrossed to the WT (Figure 1B) (Karimi-Ashtiyani et al., 2015; Kuppu et al., 2015; Maheshwari et al., 2015; Ravi and Chan, 2010). When grown, they appear normal in all respects including centromeric localization (Fig 1b) (Kuppu et al., 2020; Maheshwari et al., 2017), except *GFP-ts*, which displays somewhat stunted growth and partial male sterility. This raises several questions: when and how is GE triggered? Do the different types of CENH3-based HIs share common mechanistic features? Here, we elucidate the mechanistic basis behind CENH3-based GE during early seed development by combining genetic, molecular and cytological approaches. Using a variety of Arabidopsis HIs, we show that, in eggs of haploid inducers, the variant CENH3 is removed from the centromeres before fertilization while it persists in wild-type eggs. In GE crosses, we observed preferential CENH3 loading on WT centromeres. In the ensuing embryonic mitoses, the CENH3-depleted HI chromosomes formed defective kinetochores and missegregated frequently. In stark contrast, in crosses between HIs, CENH3-depleted centromeres undergo balanced CENH3-reloading. Mutation in a ubiquitin ligase, VIM1, increases GE efficiency. Thus, the epigenetic imbalance between parental centromeres results in parent-specific GE.

## Results

### Biased loading of GFP-ts in zygote precedes GE

In Arabidopsis, GE can be induced by fertilizing female parents expressing certain CENH3 variants (Figures 1A, and 1B) with wild-type pollen (GE cross). Among the variants, GFP-ts is an efficient HI and has the advantage of marking centromeres. Because variant CENH3s are recessive (Figure 1D,E), a plant expressing both the GFP-ts and the native wild-type CENH3 has a wild-type phenotype and does not induce GE when crossed with wild-type parent. Because of its convenient features, we employed this line in our control crosses throughout this investigation (Figure 1D). We searched for signs of GE in zygotes and in early embryonic mitoses (Figure 1B) (Ishii et al., 2015; Sanei et al., 2011; Subrahmanyam and Kasha, 1973), marking female and male chromatin with histone H2B fusion tags (Figure 1C, Methods). In both control (*CenH3;GFP-ts* X WT) and HI (*cenh3-1*;*GFP-ts* X WT) crosses, centromeric GFP-ts signals were absent in the mature haploid egg cell (Figures 1F and 1G). Following fertilization, centromeric GFP-ts signals were still absent in the diploid zygote (Figures 1H1,1I1, S1A-C, S1E-G,) until 19 hours after pollination (HAP). From 20-35 HAP, as zygotic G2 chromosomes condense and progress towards mitotic division (Lermontova et al., 2006), the maternally inherited GFP-ts started appearing on all 10 centromeres in the control cross (Figure 1H2), while they only appeared on 5 centromeres in the GE cross, consistent with loading on only one parental set of chromosomes in the latter (Figure 1I2). By the start of mitosis (26-36 HAP), the majority of the zygotes in both control (100%, n=38) and GE (86.4%, n=125) crosses displayed GFP-ts signals. The 10 vs. 5 patterns persisted through the first zygotic mitosis (Figures IH3-1H5, S1D vs. 1I3-1I5, S1H). No segregation abnormalities were visible in the control. In contrast, in the HI embryos, laggards and micronuclei lacking centromeric GFP-TS were observed during anaphase and post telophase cells (Figures 1I4 and 1I5). Taken together, in the control, GFP-ts displayed the expected zygotic reprogramming (Ingouff et al., 2010) of ten centromeres. In the GE cross, however, it reloaded on the five centromeres associated with properly segregating chromosomes.

Widespread seed death in the haploid induction cross could be explained by endosperm failure (Carputo et al., 1999; Zhang et al., 2016) (Figure 1C) hastened by missegregation of HI chromosomes (Ravi et al., 2014). To test this hypothesis, we examined the behavior of HI chromosomes in the endosperm. The endosperm proliferates rapidly (from ∼11 HAP) following fertilization of the central cell. In contrast, the zygote takes ∼30 HAP for its first division (Gooh et al., 2015). In the central cell nuclei, centromeric GFP-ts signals were also absent before fertilization in both control and HI lines (Figure 2F, 2G), indicating loss of GFP-ts in both female gametes. Upon completion of double-fertilization (karyogamy), the paternal chromatin can be followed by its lingering association with fluorescence of the sperm-specific histone HTR10-RFP fusion (Figures 1J-1M, S1A-B, S1E-F), which is even more pronounced in endosperm with condensed chromatin (Figures 1K, 1M and S1B, S1E-F) than in the zygote (Figures S1A and S1E). After karyogamy, in the reprogram of control crosses, maternally expressed GFP-ts was rapidly loaded on all 15 parental centromeres (84%, n=94 ovules, Figures 1J and S1A) as opposed to only 5 (90%, n=27 ovules, Figures 1L) in the GE cross. More importantly, all 5 signals were predominantly associated with distinctly marked male chromatin through the next cell division (Figures 1M, S1E and S1F). As HTR10-RFP is no longer brightly visible after the second mitosis, we tracked the later stages endosperm using *pRPS5A-H2B-tdtomato* (Maruyama et al., 2015), a constitutive marker provided by the male parent. In subsequent endosperm mitoses (20-36 HAP) of the GE cross, in addition to nuclei with 5 centromeric GFP-ts signals, we often detected nuclei with 10 or 20 brighter signals or a variable number of bright and faint signals (Figures S1G and S1H). In contrast, up to 15 signals were consistently observed in the control cross (Figures S1C and S1D). As in the embryo, rare chromosome bridges (Figure 1M, white arrowheads) and micronucleus (Figure S1H, white arrow) were found in endosperm nuclei from the GE cross.

**Figure 2:**
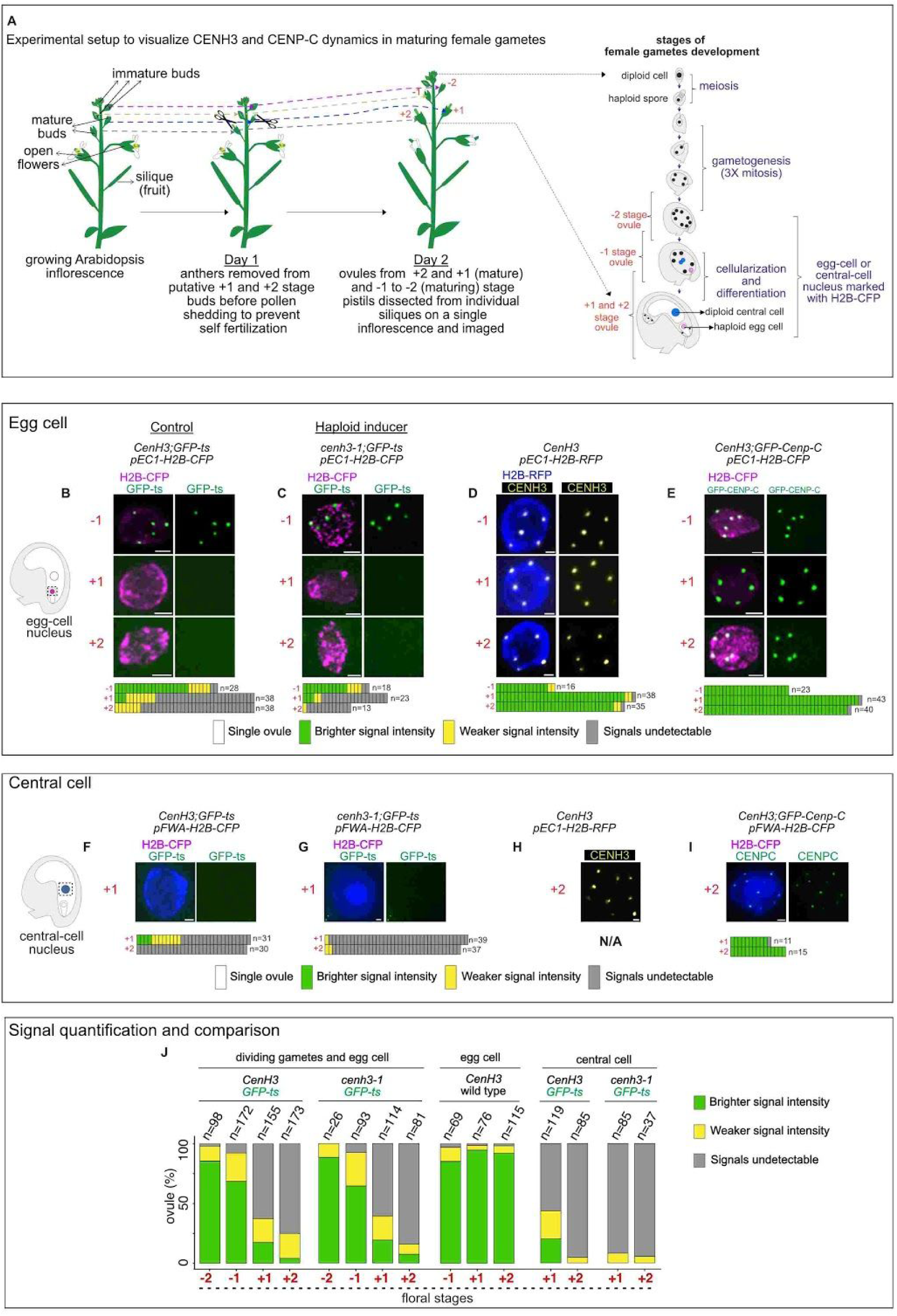
Stability of endogenous CENH3 and CENP-C. (A) Experimental setup to study the dynamics of kinetochore proteins during arabidopsis female gamete development. The left schematic indicates the region of interest in the ovule shown on the right (B-I). (B-E) Images from a single inflorescence of the respective genotypes from −1 to +2 stage ovules. (Quantification of signal dynamics in egg cells or central cells obtained from individual inflorescences are given below B-G and I). Dynamics of GFP-ts in egg-cell nuclei of control (B) and haploid inducer (C). (D) CENH3 signals (yellow, by immunostaining) and (E) GFP-CENP-C signals in the egg nuclei from the wild-type lines. GFP-ts signals in differentiated central-cell nuclei of control (F) and haploid-inducer (G). CENH3 signals (yellow, by immunostaining) in the presumptive central-cell nucleus (H) and GFP-CENP-C signals in the central-cell nucleus (I). (J) GFP-ts and CENH3 signal quantification in egg and central cells from multiple inflorescences of control, haploid inducer and wild-type lines. “N/A” not available. Scale bar=1 µm.

Quantification of GFP-ts reloading timing in both the embryo and endosperm (Figure 1SI and ISJ) revealed a similar dynamics in both control and GE crosses. These observations appeared to confirm zygotic reprogramming of CENH3 (Ingouff et al., 2010). During GE, however, GFP-ts localizes to five centromeres, consistent with paternal bias and failure to reload on the centromeres inherited from HI parent (concluded from observations on endosperm; Figure 1 and S1). This raised two questions: first, if CENH3 vacates centromeres as proposed (Ingouff et al., 2010), how is centromere identity preserved in the egg cell and central cell? Second, what determines biased loading in the GE cross?

### Native CENH3 and GFP-CENP-C display stable inheritance during development of female gametes and zygote

In embryos from control crosses, where the female parent expressed both GFP-ts and CENH3 (Figure 1), the maternal chromosomes retained and propagated centromeric identity, even if centromeric GFP-ts was absent in mature eggs. Could GFP-ts be removed selectively, while CENH3 persisted? In addition to eviction and reloading of CENH3-GFP variant (Ingouff et al., 2010), previous analysis of male meiosis suggested this possibility (Lermontova et al., 2011a; Ravi et al., 2011). To explore the stability of GFP-ts and wild-type CENH3, we compared immature (flower stage -2,-1) and mature (flower stage +1,+2,) ovules (Borg et al., 2020; Schneitz et al., 1995; Smyth et al., 1990) bracketing the normal time duration of self-fertilization (Figure 2A). In both HI and control lines, while the five and ten centromeric GFP-ts signals disappeared respectively from a majority of egg and central cell after stage -1 (Figures 2B, 2C, 2F and 2G), wild-type CENH3 (detected by whole-mount immunostaining of ovules) persisted in both egg (Figure 2D) and central cells of +1 (data not shown), +2 (Figure 2H). In comparison to the stable presence of CENH3 in the mature female gametes from wild-type, GFP-ts appeared to be actively removed from the centromeres in the mature gametes from both control and HI lines (Figure 2J). When the HI was the male, paternally inherited GFP-ts was still visible immediately after fertilization (around 9HAP), but disappeared thereafter until reloaded ∼25 HAP (Figure S2B). However, in endosperm, paternal GFP-ts readily marked all 15 centromeres from fertilization onward through development (Figure S2B). The association of GFP tagging with removal from chromatin is peculiar to CENH3, because analogous fusions with CENP-C and Nuf-2 were retained in the kinetochore (Figures 2E, 2I and S2C-G).

In summary, in the mature egg and central cell, wild-type CENH3 was retained in the centromeres while GFP-ts was evicted. Mechanisms that selectively remove altered CENH3 and kinetochore proteins exist both in male (Figure 2H) and female gametes (Ingouff et al., 2010). When transmitted by pollen, GFP-ts was removed from the zygote. While altered GFP-ts is actively removed in egg, central cell and zygote, wild-type CENH3 and other GFP-tagged key kinetochore components, such as GFP-CENP-C or NUF2-GFP, are retained through fertilization.

### Interploidy GE crosses confirm depletion of GFP-ts from HI parent chromosomes

The above data suggested that centromeres inherited from one parent, presumably the HI, are incompetent for CENH3 loading. To test this hypothesis, we marked the genome of origin using parents that differed in ploidy. Interploidy crosses, such as 4x(tetraploid) X 2x(diploid) and the reciprocal cross, are possible in Arabidopsis (Henry et al., 2005; Scott et al., 1998), and haploid induction has been demonstrated in both cases (Marimuthu et al., 2011; Ravi et al., 2014). In the 3x embryos produced by a control interploidy cross (2x *CENH3(+/+);GFP-ts* X 4x WT), most of the cells exhibited 15 or 14 centromeric GFP-ts signals as expected (Figures 3A). The cells of 3x embryos from the GE cross (2x *cenh3-1;GFP-ts* X 4x WT) carried 15 chromosomes, 5 maternal and 10 paternal. If the maternal GFP-ts was associated only with the wild-type, paternal centromeres, we should observe 10 signals. Indeed, we found that a majority of nuclei displayed 10 brighter signals, often with an additional 5 or fewer faint signals (Figure 3B). Conversely, in 3x embryos from the reciprocal ploidy GE cross (4x *cenh3-1;GFPts* X 2x WT) (Figure 3C), most nuclei exhibited five brighter signals, along with 7 to 11 fainter signals. Furthermore, in 4x *CenH3 (+/-);GFP-ts* x 2x WT cross, most of the nuclei (n=10/11 embryos) exhibited the expected 15 signals, except for one embryo, in which every nucleus exhibited five bright signals along with 9 or 10 fainter signals (Figure 3D). We hypothesized that the embryo with 5 brighter signals inherited two null *cenh3-1* alleles (see test in Figure 8B), and the rest inherited at least one wild-type allele. Analysis of the GFP-ts signal number and pattern per cell (Figures 3A-D) and quantification of the GFP-ts signal intensity in a subset of the nuclei (Figure 3E) from the four crosses described above resulted in the following conclusions: i) maternally inherited GFP-ts displayed a clear bias in localization in GE crosses only. In those cases, (ii) it was predominantly found on the centromeres inherited from the WT parent. (iii) The appearance of faint centromeric GFP-ts signals in 2-4 cell stages (as found in later stages of endosperm Figure S1H) was consistent with slow, progressive reloading of kinetochore components on revenant maternal centromeres, as documented below in Figure 7E.

**Figure 3:**
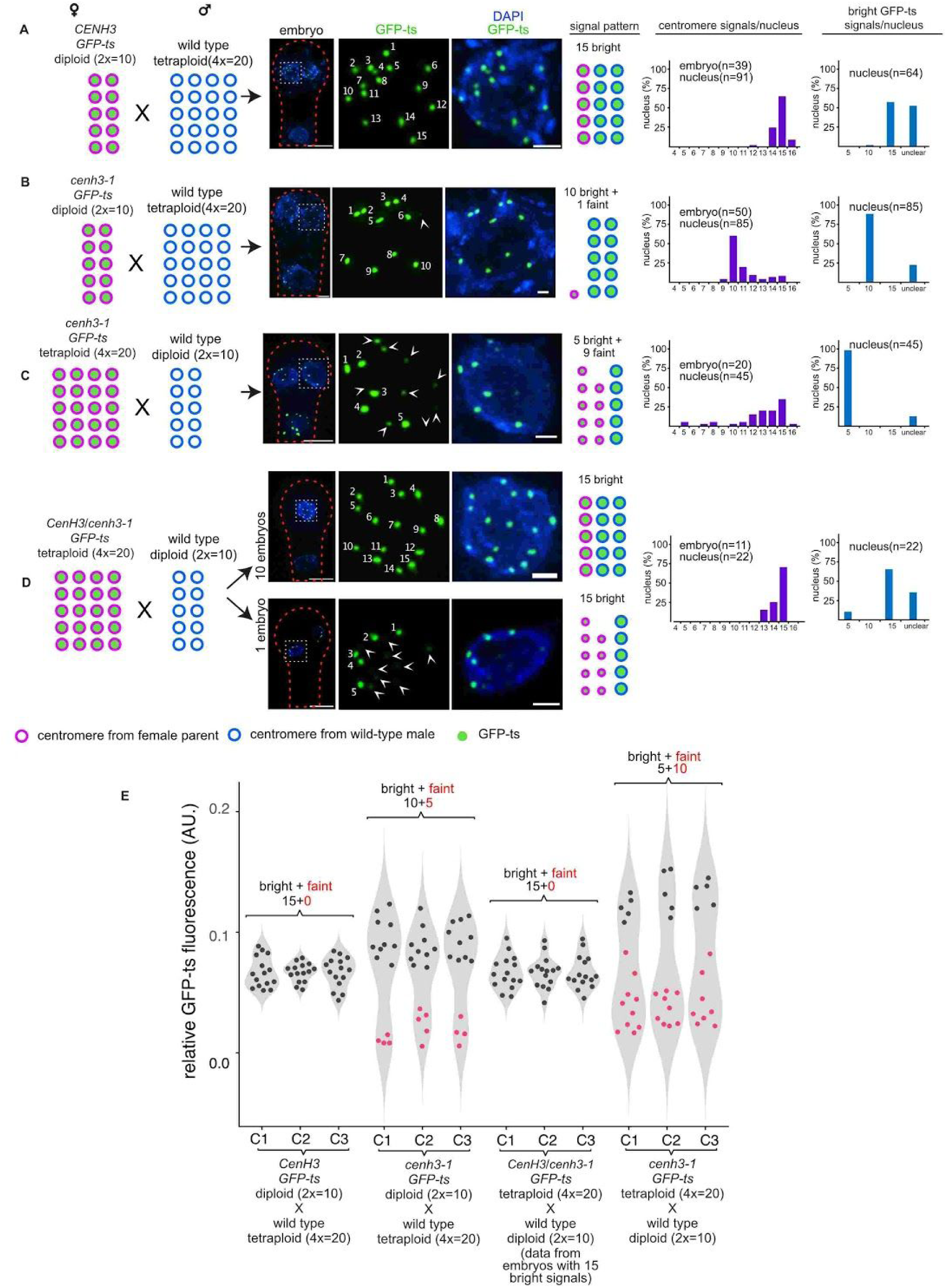
Interploidy GE crosses confirm depletion of GFP-ts from HI parent chromosomes. Tetraploid **(A, B)** and diploid **(C, D)** Arabidopsis wild-type strains were crossed as males with diploid **(A, B)** or tetraploid **(C,D)** females expressing GFP-ts in *CenH3+/+* (**A**) *cenh3*-1 -/- (**B,C**) and *CenH3+/-* **(D)** background. Arrowheads mark the faint GFP-ts signals. (**E)** Relative fluorescence intensity of centromeric GFP-ts signals. Fainter signals from the HI-crosses are highlighted in red circles. Each column represents normalized intensity in arbitrary units (AU.) within a single nucleus recorded from three different cells (C1-C3) for each cross. Scale bar=5 µm for embryo and 1µm for individual nucleus.

### HI centromeres sustain partial centromeric identity

If centromeres in the HI female gametes (Figures 1, 2, and 3) lose their identity in an outcross, how do the HI chromosomes maintain stability during self-pollination? We hypothesized that loss of identity is partial since haploid induction never exceeded 46%. To document how centromeres may regain in functionality after the zygotic stage, we first examined those generated by self-pollination in WT (Figures S3A and S3I), *CENH3;GFP-ts* (Figures 4A and S3C), *cenh3-1;GFP-CenH3* (Figures S3B and S3J), and *cenh3-1;GFP-ts* (Figures 4B and S3D). In all types of 2 to 4-cell embryos, the modal value for the centromeric signals was 10, as expected. Centromere signal intensity, displayed similar variation (1.4-1.8 v.s 1.5-1.8 fold) and relative intensity distribution (0.06-0.13 v.s. 0.07-0.14 AU.) per centromere in WT and GFP-ts, respectively. Embryos from *CENH3;GFP-ts* X WT (control cross) displayed the expected 10 signals (Figures 1C and S3E) with intensity distribution profiles similar to the self-pollinated controls.

**Figure 4:**
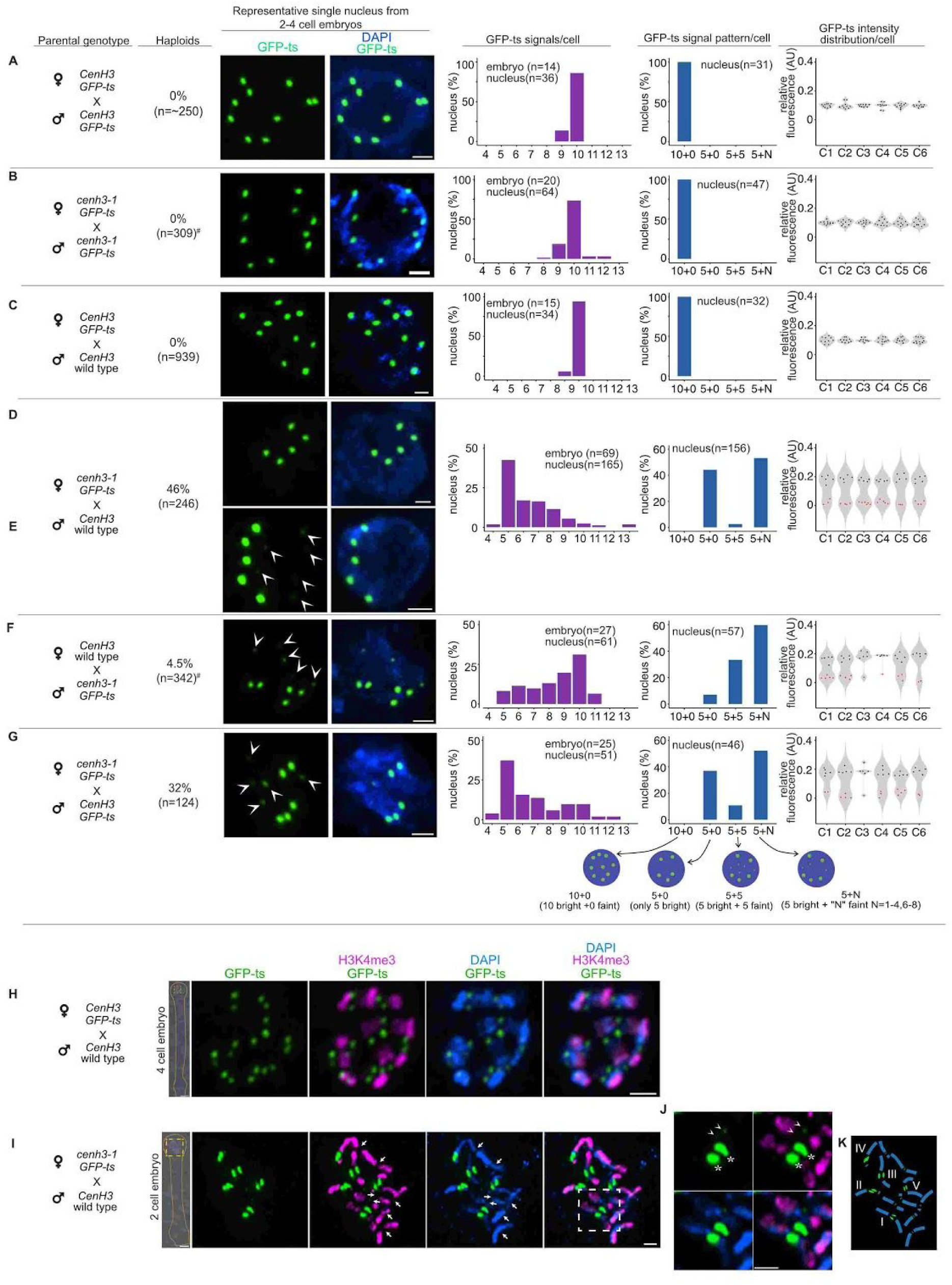
Uniparental loading of GFP-ts on centromeres of 2- to 4-cell stage hybrid embryos during GE. (A-C) GFP-ts display ten strong centromeric signals early embryonic nuclei in control crosses, while only 5 strong signals are visible in GE crosses (D-G). In some cases, up to six weak signals (arrowheads) are also visible (E-G), indicating sub optimal loading one to one parental centromere set. “n”: number of nuclei or embryo scored. The green channel is enhanced to reveal the fainter signals (E-G). Each column of the violin plots indicates relative GFP-ts intensity from six cells (C1-C6) with fainter GFP-ts signals highlighted in red circles. (H,I) whole embryos are shown on the left side and the highlighted nuclei with the yellow dotted line shown on the right. (J) selected chromosomes are highlighted in I with white dotted box. Prometaphase chromosomes demonstrate biased GFP-ts localization in HI, but normal H3K4me3 localization away from the pericentromere and centromere. Arrows mark maternal chromosomes; arrowheads indicate faint GFP-ts signals; “*” indicates bright signals. (K) Inferred wild-type karyotype and location of brighter GFP-ts signals of the marqueed nucleus in I. “#” Data from (Ravi and Chan, 2010; Tan et al., 2015). Scale bar = 1um for single nucleus and 5um for whole embryo images.

In contrast to results in the self-pollination and control cross, 2 to 4-cell embryos from the GE cross only displayed five bright nuclear GFP-ts signals. However, differently than in the zygote, these nuclei displayed 1 to 8 fainter GFP-ts signals along with the 5 brighter signals (n=165 nuclei, 69 embryos). Both bright and faint signals were distributed randomly across the different z-sections in a nucleus, thus ruling out photobleaching while capturing images (data not shown). The brightest signal was 13.6 to 33.9 fold stronger than the weakest signal within each nucleus (Figures 4D, 4E and S3F). Though we found 1.1 to 1.5 fold intensity variation among the brighter signals, the fainter signals displayed even wider variability (1.4 to 5.3). Collectively, the bright and weak GFP-ts centromeric signals formed the following three recognizable qualitative patterns (strong+weak) within each nucleus: (i) 5+0, (ii) 5+5, and (iii) 5+N (where N=1-4,6-8). Overall, 95% of the nuclei (n=165 from 69 embryos) displayed at least five bright signals compared to 10 in all of the controls. Of those 69 embryos, 17% displayed only the 5+0 pattern on all observed cells, whereas the rest displayed combinations of all three patterns.

To further interpret the GFP-ts distribution pattern, we imaged cells in the embryo with condensed chromosomes. As expected for prometaphase, embryo cells from control crosses displayed ten pairs of bright signals (Figures 4H and S3K), while those from GE crosses displayed five pairs of bright signals (Figures 4I and S3L). Additionally in the same cell, paired but faint signals were visible on multiple additional chromosomes (up to 7) (Figures 4J and 4K). The appearance of faint signals in 2 to 4-cell embryos was intriguing as these were not readily detected in the earlier zygotic stage. The appearance of mixed patterns (5+0, 5+5, and 5+N) of centromeric GFP-ts signals in an embryo, and comparison to controls, suggested that the faint signals detected in 2 to 4-cell embryos are not due to optical distortion in embryo whole-mounts. Instead, they correspond to gradual reloading of GFP-ts centromeres after the zygotic transition.

In the reciprocal cross (WT X *GFP-ts*), which generates 10 fold fewer haploids than *GFP-ts* X WT (Ravi and Chan, 2010), cells with only 5 bright centromere signals were less common: 7% 5+0 pattern vs 44% for *GFP-ts* X WT (p<.0001, 2-sample z-test), suggesting that reloading of HI centromeres was more efficient in crosses with diminished GE potential (Figures 4F, S3G). Biparental provision of GFP-ts (*cenh3-/-*;*GFP-ts* X *CENH3; GFP-ts*, 32% haploid progeny, n=124, Figure 4G, S3H) did not alter the GFP signal patterns, which remained similar to GFP-ts X WT. Additionally we tested the effect of the mcherry fluorescent tag using *GFP-ts* x *mcherry-ts;CENH3* cross. As observed for GFP-ts, the paternal mCherry tag did not alter the outcome and embryos displayed 5 bright GFP-ts signals as in *GFP-ts* X *CENH3:GFPts,* and were colocalized with bright mCherry-ts signals (Figure S3M). These observations suggested that competitive centromeric reloading is effectively triggered by the near absence of any CENH3 on one gametic centromere set and not by the presence of altered CENH3.

Taken together, our analysis of the centromeric GFP-ts patterns in 2 to 4-cell embryos revealed the following properties of GFP-ts on *Arabidopsis* centromeres: (i) A self cross of the HI strain mimics the wild-type behavior, resulting in proper premitotic reloading of GFP-ts on all centromeres in spite of its prior removal. (ii) Mitotic condensed chromatin confirms biased GFP-ts occupancy of centromeres inherited from one parent. (iii) Male-expressed GFP-ts is evicted in the zygote and reloaded on centromeres contributed by CENH3 positive females (Figure S2B). (iv) Lower haploid yield in paternal GE crosses is correlated with increased signals on the HI-contributed centromeres (decreased 5+0 and increased 5+5 and 5+N patterns), suggesting that the lower GE efficiency of the male HI is due to higher rate of reoccupancy of HI centromeres.

### In GE crosses, wild-type and variant CENH3s localize to centromeres contributed by the wild-type parents

We wondered whether the native CENH3 contributed by the wild-type parent in a GE cross displayed the same bias as GFP-ts. To address this, we localized both CENH3 types in the 2 to 4-cell embryos by combining immunological detection of the CENH3 N-terminus (Talbert et al., 2002) with that of GFP-ts via anti-GFP antibody. Notably, the N-terminus antigens are absent in GFP-ts (Figures 1A and S4A). In the control cross, both GFP-ts and wild-type CENH3 variants colocalized on all ten centromeres, consistent with equal recognition of all ten parental centromeres (Figure 5A).

**Figure 5:**
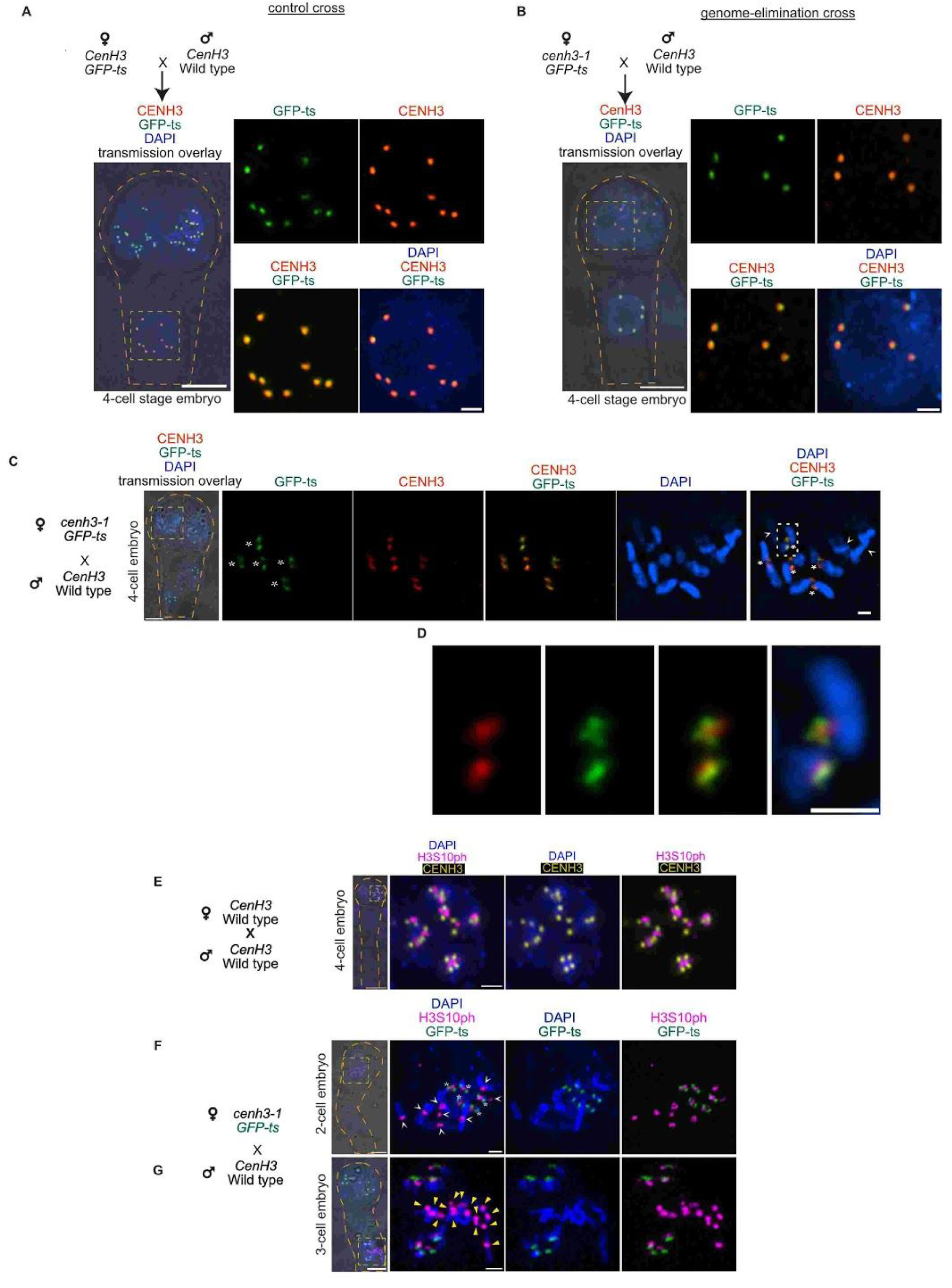
Native CENH3 and GFP tailswap-CENH3 occupy the same functional centromeres during haploid induction. (A-G) A selected nucleus (yellow dotted box) on the embryo is shown enlarged on the right. (A) Control cross illustrates colocalization of CENH3 and GFP tailswap-CENH3 on ten centromeres. (B) GE crosses display colocalization as five strong signals. (C) 3D-SIM image of prometaphase stage cells in embryos from the GE cross. (D) Enlarged condensed chromosome from image C (white dotted box). (E) At the pericentromeric region of all chromosomes, H3S10ph is noticeable as an independent domain in prometaphase stage nuclei in the wild-type embryo, and in prometaphase (F) and anaphase (G) in the embryo from the GE cross. “*” marks wild-type centromeres whereas the arrowheads mark the chromosomes lacking functional CENH3 variants. Yellow triangles (G) mark laggard chromosomes with H3S10ph signals. Scale bar = 5μm for the whole-embryo images and 1μm for the rest of the figure.

In the GE cross, both the variants colocalized on five chromosomes inherited from the wild-type parent in interphase (Figure 5B), G2 (Figure S4B), and prometaphase cells (Figure 5C). At the higher resolution provided by 3D-SIM, the two variants only overlapped partially (Figures 5C and 5D), suggesting the formation of centromeric subdomains enriched with one or the other protein, consistent with previous observations with natural or artificial combinations of CENH3 types (Ishii et al., 2015, 2020; Maheshwari et al., 2017). Often, faint GFP-ts signals are colocalized with faint CENH3 signals (Figure S3C).

Given that GE can result from crossing the CENH3 point-mutation variant M4 (Kuppu et al., 2015) with the WT, we asked if a missense mutation in CENH3 could result in similar outcomes. Consistent with previous observations, embryos from *M4 (CENH3^G83E^)* X WT cross also displayed only five centromeric CENH3 signals in the interphase nucleus and often with few faint signals as seen for *GFP-ts* X WT (Figure S4D). Imaging the anaphase stage revealed that this uniparental localization of CENH3 is specific to the leading chromatids and absent in the lagging ones (Figure S6B). A control point mutation (M47) in another highly conserved HFD residue, G173E, does not act as an HI (Kuppu et al., 2015). As expected, the embryos from this cross did not display biased loading of CENH3 when crossed to the wild-type male, displayed 10 bright CENH3 signals and normal segregation (Figures S4E and S6C). We concluded that all tested CENH3s, native or GE-inducing, display similar localization bias in GE crosses. The concordant response of different CENH3s confirms that only the wild-type centromeres are competent during the very early embryonic mitosis, and able to assemble the functional kinetochores (Figure 3). In addition, it suggests that the CENH3^G83E^ variant, like GFP-ts, is removed during egg maturation.

### In GE crosses, typical CEN chromatin states persist on defective chromosomes

We wondered whether the loss of CENH3 during GE was associated with remodeling of the stereotypical chromatin states of the centromere and pericentromeric regions. Phosphorylation of H3 Serine 10 (H3S10ph) is found in many plants on pericentric chromatin but restricted to condensed chromosomes (Houben et al., 1999, 2007; Kaszás and Cande, 2000; Shibata and Murata, 2004). On the other hand, H3K4me3 marks the euchromatic region but is excluded from the centromere proper (Sullivan and Karpen, 2004; Zhang et al., 2008). In the pro-metaphase stage of embryos from the GE cross, chromosomes inherited from both parents displayed the H3S10ph signals (Figure 5F) and were similar to WT (Figure 5E). Even after sister chromatid cohesion resolved at anaphase, we found H3S10ph on leading sister chromatids with functional centromeres, as well as on the lagging chromatids, which presumably lack centromere function (Figure 5G). Interestingly, H3S10ph signal intensity on the laggard chromosomes appeared stronger than the leading chromosomes. Importantly, detection of >10 (14 H3S10ph signals on the laggard chromosomes in Figure 5G) H3S10ph signals suggested the presence of dicentric chromosomes (Shibata and Murata, 2004) in a subset of those laggards. Similarly, on chromosomes inherited from both wild-type and GFP-ts parents, the euchromatic specific H3K4me3 was clearly excluded in pericentric and centromeric region but strongly stained the euchromatic arms (Figure 4I and 4J). The H3K9me2, a heterochromatic mark, was also found on both leading and lagging chromatids although precise boundaries could not be compared (Figure S6B).

In conclusion, because normal centromeric patterns of H3 modification persisted after differential CENH3 loading and during centromeric failure, they are unlikely to underlie chromosome missegregation and loss.

### A single parental chromosome set assembled functional kinetochores

We reasoned that, if the biased loading of GFP-ts is a cause of uniparental centromere dysfunction, this outcome should be reflected in a kinetochore defect. To demonstrate this, we examined CENP-C, a pivotal inner-kinetochore protein acting downstream of CENP-A and NUF2, a component of the outer kinetochore complex that directly interacts with the spindle-microtubules (Cheeseman and Desai, 2008; Klare et al., 2015; Musacchio and Desai, 2017; Ogura et al., 2004; Screpanti et al., 2011; Tomkiel et al., 1994; Wei et al., 2005). Both fusion proteins produced ten fluorescent signals in somatic nuclei (Figures 3A and 3B). When paternally contributed in control crosses, both marked all ten parental centromeres equally in 2-4-cell stage embryos (wild-type female: Figures 3C and 3D) and colocalized with maternal GFP-ts (*CENH3;GFP-ts* female: Figures 3E and 3G), and matched it in intensity.

In the corresponding GE crosses, similar to GFP-ts signals, both RFP-CENP-C and NUF2-RFP signals (Figures 3F and 3H) displayed wide numerical variation and fell into distinct bright and faint qualitative categories whose intensity differed, respectively, 9 to 60 fold and 5 to 65 fold. Like GFP-ts, both RFP-CENP-C and NUF2-RFP followed 5+0, 5+5 and 5+N (where N=1-4 and 6-8) patterns. Interestingly, in both crosses, bright RFP signals colocalized with the bright GFP-ts signals. Singleton faint signals were rarely observed (Figures 6F, 6H, asterisk marks). This biased loading pattern was also apparent in later stage embryos (3-5 days after pollination, data not shown). We concluded that in GE crosses, only the wild-type parental set of centromeres loaded both GFP-ts and wild-type CENH3 (Figure 5), and assembled functional kinetochores.

**Figure 6.**
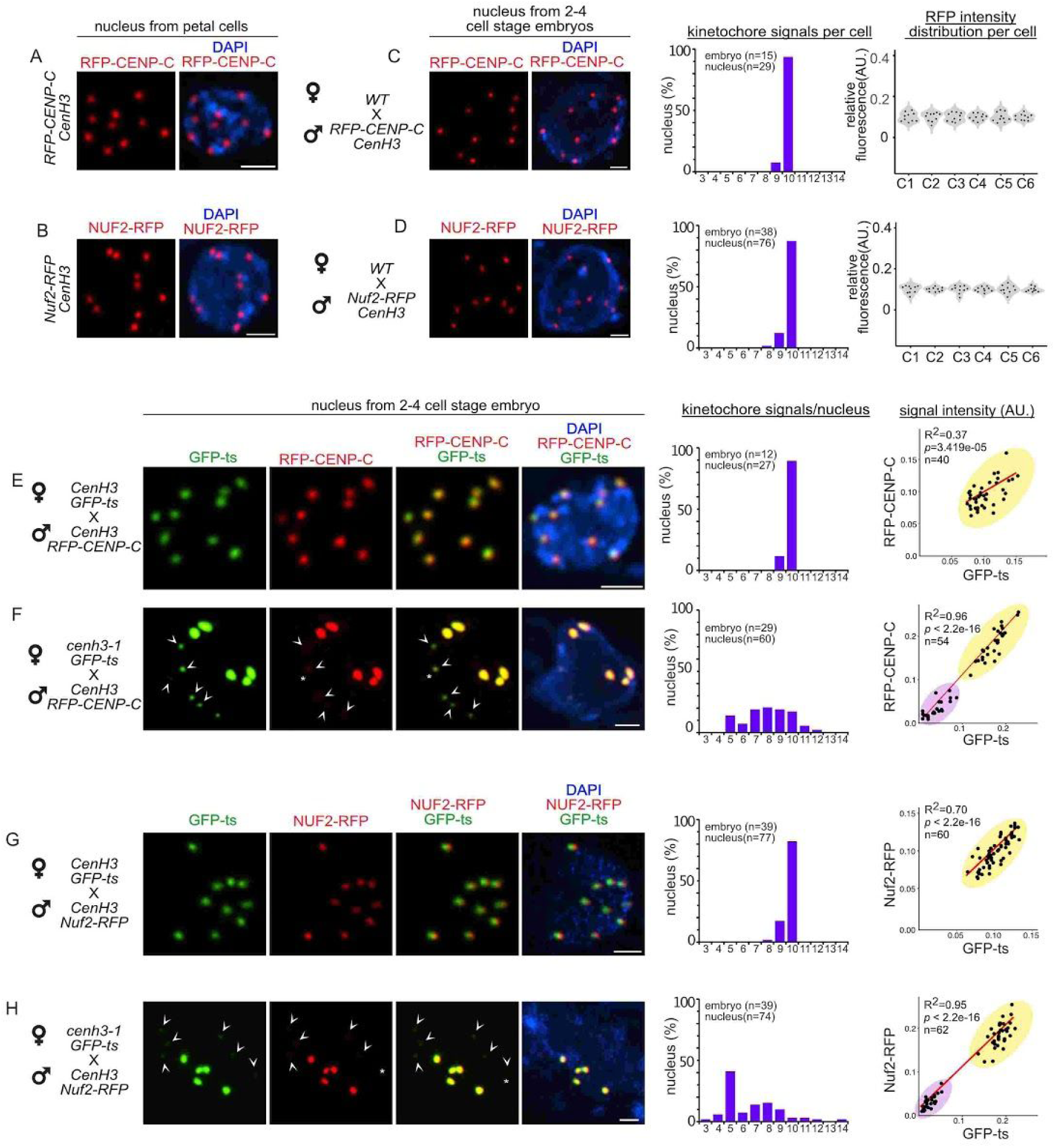
Uniparental assembly of functional kinetochores in embryos undergoing GE. (A,B) Localization of RFP tagged CENP-C, and NUF2 in petal cells. (C-H) Interphase nuclei images from 2 to 4-cell stage embryos. Localization of paternal RFP tagged CENP-C (C), and NUF2 (D) on all 10 parental kinetochores in control cross with wild-type females. Comparison of GFP-ts, CENP-C, and NUF2 colocalization in interphase nuclei from 2 to 4-cell stage embryos in control (E,G) and GE crosses (F,H). White arrowheads and “*” in F and H, respectively, mark faint signals and singleton GFP or RFP signals. Bar graphs, violin plots and correlation plots represent the kinetochore numbers and relative fluorescent intensity values for the corresponding genotypes shown on the left. Each column of the violin plots indicates relative RFP intensity from six cells (C1-C6). Scale bar=1um.

**Figure 7.**
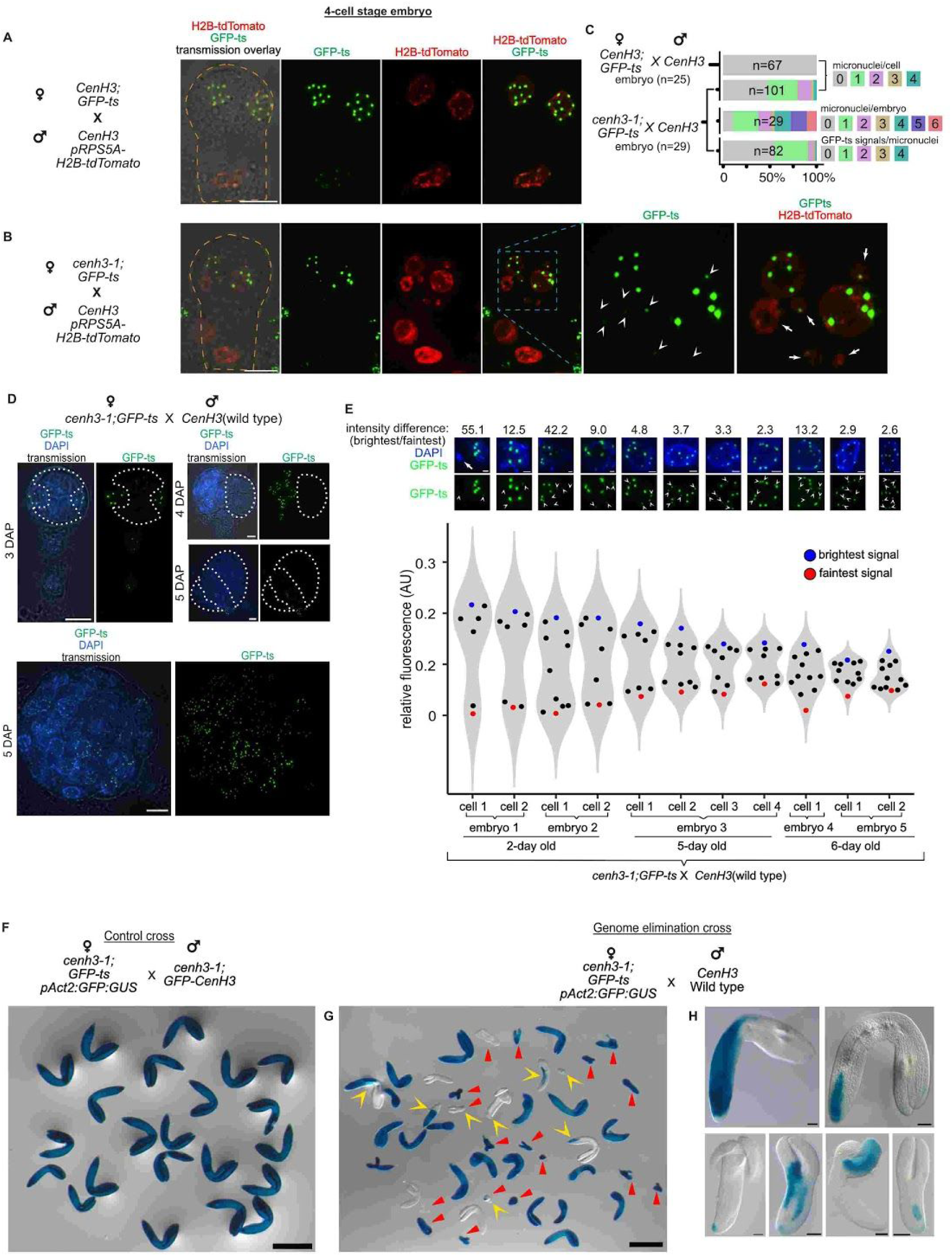
Fate of HI chromosomes in developing embryos. Early embryonic divisions in control (A) and GE cross (B). (B, right) Micronuclei (white arrows) containing weakly labeled HI chromosomes (white arrowheads). (C) The bar graph depicts quantitative analysis of micronuclei and GFP-ts signals in control and HI crosses represented in figures A and B. (D) Chimeric distribution of GFP-ts signals in early embryos indicates elimination of the GFP-ts-encoding chromosome (HI genome) in cells lacking signal or displaying fainter signals (highlighted with dotted white line). (E) Resiliency of HI centromeres demonstrated by progressive coalescence of bright and faint GFP-ts fluorescence intensity clusters and by the decrease in the maximum-minimum difference (white arrowhead: weaker signals; white arrow: micronuclei). GUS histochemical stain locates uniformly in two-week-old control embryos (F). (G, H) GE cross. (G) GUS staining chimerism indicates (yellow arrowheads) frequent incomplete GE during embryo development. Smaller embryos (red triangles on a sub-set) likely result from severe aneuploidy or defective endosperm. (H) chimeric embryos at a higher magnification. Scale bar = 5μm for A, B and D, 1μm for E; 500μm for F and G; 50μm for H.

**Figure 8:**
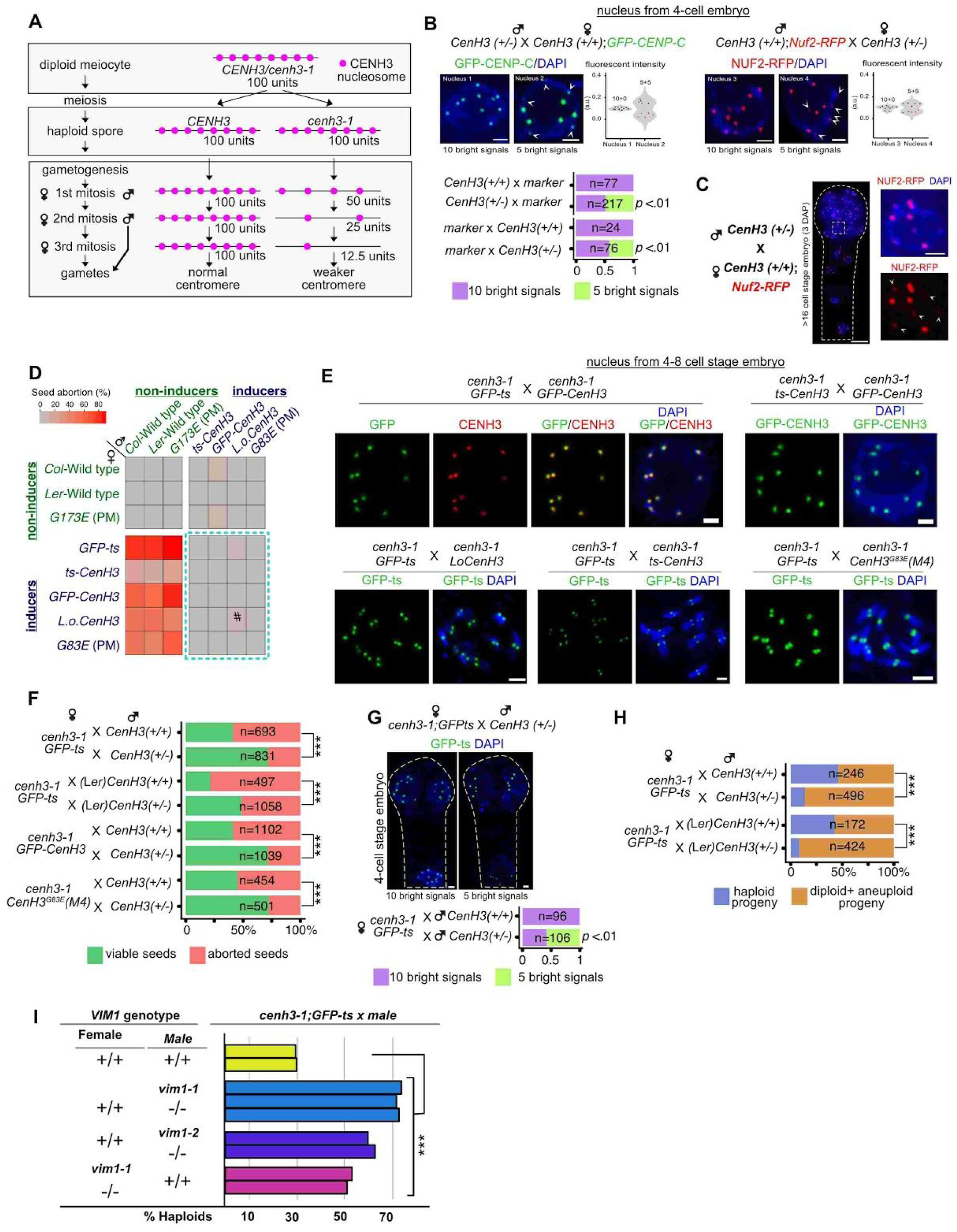
CENH3 dependent and independent factors alter seed death and GE efficiency. (A) Hypothetical dilution of centromeric CENH3 during gametogenesis following segregation of wild-type and null allele in meiosis. (B) Patterns of kinetochore signal intensity in embryos generated by crossing *CENH3(+/-)* parent with lines expressing kinetochore markers. White arrowheads indicate fainter kinetochore signals. Bottom, counts for embryos shown in B. Violin plots display kinetochore signal intensity from the images on the right. Red circles represent faint signals. (C) Persistence of bright and faint kinetochore in >16 cell stage embryos. (D) Heat map of seed abortion (a proxy for GE frequency) when intercrossing non-inducer and HI lines. ^#^: Data from (Maheshwari et al., 2015) (E) Normal, 10-chromosome loading of CENH3-GFP or GFP-ts in crosses between GFP-ts and other HI lines (interphase or prometaphase stage nuclei). (F) Proportion of aborted seeds upon pollinating various HI lines with *CenH3* (+/-) males. (G) Embryos from *GFP-ts* X *Cenh3(+/-)* with two patterns of GFP-ts signal. Bottom, counts for embryos in G. (H). Bar graph displays GE induction efficiency when pollinating GFP-ts with *CenH3* (+/-) males. “n” indicates the number of embryos or progeny quantified. (I) Bar graph showing the effect of null alleles of *VIM1* on *GFP-ts* haploid induction efficiency. Scale bar=1μm.

We asked whether the other CENH3 variants (Figure 1A) displayed loading dynamics similar to GFP-ts. We pollinated different haploid inducers(*GFP-CENH3*, *tailswap-CENH3*, *LoCENH3*, *ZmCENH3*, *M4(CenH3^G83E^)*, *A86V(cenh3-2)* and a non-haploid inducer *M47 (CenH3^G173E^) (Kuppu et al., 2015; Maheshwari et al., 2015; Ravi and Chan, 2010; Ravi et al., 2010)* with male plants expressing either NUF2-RFP or NUF2-GFP or GFP-CENP-C, but carrying wild-type CENH3. Embryos from all GE crosses displayed bright and faint kinetochore signals whose count and pattern were comparable to those of GE crosses with GFP-ts (Figure 4D). At least 55-93% nuclei in all GE crosses displayed five-strong kinetochores as 5+0, 5+5 and 5+N patterns with genotype specific variations (Figures S5A-S5F). A 5+0 pattern signifies strong uniparental loss of centromere identity, whereas the 5+5 and 5+N patterns mark the potential recovery of weak centromeres. Except for *A86V (cenh3-2),* embryos from all crosses 3-42% of observed nuclei displayed a 5+0 pattern. In contrast, when the HI was male, the 5+0 pattern was markedly absent and the 5+5 pattern were significantly more frequent (9 to 63% higher, bar graph in Figures S5A-S5F). In contrast, the non-HI-inducing transgenic variant CENH3^G173E^ assembled kinetochores that had similar intensity on all ten parental centromeres. Here, a majority of the observed nuclei displayed the 10+0 pattern (80%, n=51, in the G173E X marker cross, and 87%, n=47, in the marker X G173E cross, Figure S5G).

In summary, loss of centromere identity on HI centromeres results in defective kinetochore assembly across all tested CENH3 variants, indicating a common mechanism for GE. Further, the 5+0 kinetochore pattern is strongly correlated with GE efficiency. When the HI was the pollinator, the resulting embryos displayed predominantly the 5+5 pattern associated with improved recovery of centromeres inherited from the HI variants.

### Chromosomes with CENH3-depleted centromeres missegregate and partition into micronuclei

Centromere inactivity results in laggard chromosomes that partition into micronuclei (Tan et al., 2015), consistent with genome instability (Fujiwara et al., 1997; Gernand et al., 2005; Gibeaux et al., 2018; Ishii et al., 2010, 2016; Leibowitz et al., 2015; Ly et al., 2017; Sanei et al., 2011). To determine if chromosomes in micronuclei carried defective centromeres, we compared centromere states of chromosomes in different nuclear structures. In control crosses (Figure 7A and 7C), chromatin labeled with paternal H2B-tdTomato (Maruyama et al., 2015) was always associated with normal nuclei (micronuclei were absent; n=67 cells from 25 embryos), and displayed 9-10 distinct GFP-ts signals, as expected. In GE crosses, laggards were observed starting with the first (Figure 1I4) or second (Figure S4C) zygotic mitosis. By staining the euchromatin marker H3K4me3 (Figure 4I), we determined that laggards were predominantly lacking GFP-ts signal in the centromere (Figure S6A). In some cases, chromatin bridges devoid of GFP-ts signals were stretched between two poles (Figures 1I4 and S6A) suggestive of DNA breaks. These aberrant structures were not detected in control crosses. Similarly, laggard chromosomes were associated with the HI variant (CENH3^G83E^), but not with the non-inducer variant (CENH3^G173E^) (Figures S6B and S6C). After the second embryonic mitosis in GE crosses (n=101 cells from 29 embryos), the H2B-tdTomato chromatin was associated with two types of nuclei (Figures 7B and 7C): i) Regularly-sized nuclei, present one per cell, with five strong GFP-ts signals in 88% of cells (n=101). ii) 90% (n=29) of the embryos had at least one cell with 1-4 micronuclei of variable size. Up to 59% of them had more than one cell with micronuclei; (iii) a subset of micronuclei (45%, n=82) had one to four faint GFP-ts signals (Figures 7B and 7C). Finally, the presence of micronuclei (90% positive embryos, n=30) was strongly associated with seed death in the GFP-ts x WT cross (Figure 8D; 80%, n=602 seeds). Micronuclei of variable size were observed even in later stages of embryo development (Figures S6D and S6E). In summary, the inability to load any CENH3 on the centromeres of HI chromosomes led to failure of kinetochore assembly, missegregation of laggards, and frequent partition within micronuclei.

### Alternative fates of chromosomes with CENH3-depleted centromeres

Multiple observations suggest that centromeres can replenish centromeric-CENH3 after its depletion (Gassmann et al., 2012; Mitra et al., 2020; Raychaudhuri et al., 2012). The uniform underperformance of HI chromosomes in early embryos together with the production of diploid and aneuploid progeny from a GE cross, indicate that centromere function of HI chromosomes recovered during embryo development. Supporting this notion, HI chromosomes carrying faint GFP-ts segregated normally (Figures S6A and S6E). Further, comparing embryos in the 2-6 DAP window, we observed progressive coalescence of high and low GFP-ts signals toward uniformity (Figure 7D and 7E). If GE is stochastic and certain cells regain HI centromeric competence, chimeric sectors should be frequent in developing embryos. To test this hypothesis we examined GFP-ts signals in 3-5 days old embryos finding frequent chimerism (Figures 7D, S6F). To visualize the pattern of GE in later stage embryos, we used the histological marker GUS provided as a transgene in the HI genome. At 14 DAP, when embryos have established their embryonic axis, embryos in the control cross displayed uniform development and staining (Figure 7F), while embryos from GE cross varied widely in development and staining pattern (Figure 7G and 7H). Later, chimerism was also common in seedlings as variable size and pattern stained sectors that retained the HI chromosome carrying GUS (Figure S6G). Overall, the shoot apical meristem state reflected the composition of progeny upon germination, which encompass diploid, aneuploids, and haploids (Tan et al., 2015). Taken together, these results indicate that if HI chromosomes escape early missegregation, their centromeres compete better and better for CENH3 loading.

### Dilution of CENH3 nucleosomes mimics CENH3-dependent HI

Induced variation in Cid (a.k.a. CENP-A) level by over-expression or RNAi-based depletion in Drosophila sperms, proportionately affected the centromere strength in the next generation (Raychaudhuri et al., 2012). Removal of altered CENH3 may lower the density of CENH3 nucleosomes below a critical threshold diminishing the competitive potential of the affected centromeres. We tested the effect of depletion using plants heterozygous for the *cenh3-1* knockout mutation (+/-). Following meiosis, the haploid spores undergo 3 mitoses in the female and 2 mitoses in the male to generate gametes. Those gametes inheriting the null-allele lack continued CENH3 protein provision and should undergo progressive depletion (Figure 8A). In spite of CENH3 essentiality, the null allele (*atcenh3-1*) is transmitted efficiently when inherited from a heterozygous parent (Ravi et al., 2010), in contrast to transmission defects observed in maize *cenh3* null gametes (Wang et al., 2021).

When a *CenH3/cenh3-1* Arabidopsis was crossed to the *CenH3(+/+)* carrying kinetochore marker in either direction, half of the embryos displayed biased kinetochore signal intensity (5 strong + up to 5 weak signals) and while the remaining half displayed 10 bright signals (Figure 8B). This pattern was retained in the later stages of embryo development (Figure 8C). In contrast, all embryos from *CenH3(+/+)* parents generated embryos with 10 bright signals (bar graph, Figure 8B). More importantly, screening of the progeny from *CenH3/cenh3-1* x *CenH3(+/+)* (female listed first) identified 4 haploids/956 progeny or 0.83% of zygotes formed by (-) eggs. This highlights the importance of threshold quantity of CENH3 in centromere function and demonstrates that haploids can be induced without altering CENH3, but by simply diluting its wild-type form. Wang et al. demonstrated HI in maize using a similar approach (Wang et al., 2021).

### Factors affecting the frequency of HI

We searched for factors that affect CENH3-mediated HI. We used seed death to quantify GE efficiency (Kuppu et al., 2020; Ravi and Chan, 2010). Expanding on our previous observations (Marimuthu et al., 2011), the best suppressors of the GFP-ts HI were found to be another CENH3-based HI, including those based on fusion proteins, point mutations and diverged CENH3 (Figures 7F and 8D). A majority of the HI X HI cross generated only a background level of seed-death. In addition to suppressing seed death, most embryos from *GFP-ts* x other HIs displayed a clear 10+0 pattern (Figure 8E), a significant deviation from the GFP-ts X WT cross (Figure 4D), thus providing a visual mark of functional recovery of centromeres inherited from GFP-ts and other HIs. Consistent with these observations, even the male gametes inheriting the *cenh3-1* null allele (from *Cenh3/cenh3-1* plants) and carrying CENH3-diluted centromeres were good suppressors of seed death reducing it by 30% in three different HIs (Figure 8F). A second null allele (*cenh3-3*) generated using CRISPR-CAS9 in the *Ler* accession, acted similarly (Figure 8F). Remarkably, suppression of seed death by *cenh3*-null gametes was directly reflected in the centromeric GFP-ts signal pattern in the GE cross: in the GFP-ts x *CenH3*/*cenh3-1* cross, half of the embryos displayed ten bright signals (Figure 8G), a significant deviation from *GFP-ts* X WT (Figure 4D). Corroborating these observations, the *GFP-ts x CenH3 (+/-)* cross also generated 30% fewer haploids (Figure 8H). Thus, the *cenh3*-null gametes with reduced centromeric strength rescued both lethality and GFP-ts localization, probably by matching the female’s CENH3-depleted centromeres as observed in a HI X HI cross (Figure 8D).

In a genetic screen employing natural variation, two different null mutants of *VIM1* acted as enhancers of GE when contributed by either parent (Figure 8I). VIM1 is a homolog of the yeast E3 ubiquitin ligase PSH1 (Hewawasam et al., 2010; Kraft et al., 2008; Ranjitkar et al., 2010), which regulates stability of yeast centromeric histone CSE4. In addition, mutations of *VIM1* affect DNA methylation and CENH3 density at the centromeres (Woo et al., 2007). They increased GE when contributed by either parent, suggesting that a critically low level of VIM1 in the zygote engenders centromeric failure. The strong effect of these modifiers indicates that the ubiquitination pathway, directly or indirectly, affects CENH3 stability.

## Discussion

Explanations for GE in interspecific crosses proposed several mechanisms (Ishii et al., 2016), including uniparental centromere inactivation, potentially driven by defective CENH3 localization on centromeres of the HI parent (Sanei et al., 2011). CENH3-mediated GE in *A. thaliana* mimics the genetic consequences of interspecific crosses in a tractable intraspecific isogenic system, which we employ here to address the responsible mechanism leading to several conclusions.

GFP-ts is selectively removed from centromeric chromatin during egg maturation while wild-type CENH3 persists. While the peculiar behavior of the GFP-ts may result from its extensive alteration (Figure 1A), we documented corresponding, but less severe effects in a point mutant, the G83E variant and other tested HIs. GE efficiency varies according to the structure of the variant CENH3 forming an allelic series consistent with varying removal efficiency. Another factor affecting removal efficiency is parental origin: in the WT x male HI cross, lower efficiency of removal (Figure 4, Supplemental Figures 3 and 5) is accompanied by reduction in GE rate (Ravi and Chan, 2010), in many cases below detection (Kuppu et al., 2020). Therefore, it seems likely that all HI CENH3 variants act through the same mechanism in Arabidopsis (Karimi-Ashtiyani et al., 2015; Kuppu et al., 2020; Maheshwari et al., 2015). Removal may follow recognition of an altered kinetochore subunit (CENH3) by a surveillance system (Hewawasam et al., 2010; Niikura et al., 2019; Ranjitkar et al., 2010).

Our evidence demonstrates that GE results from an epigenetic conflict. In the zygote and following embryonic mitoses, depletion of CENH3 variants from the HI centromeres results in a large epigenetic imbalance with wild-type centromeres, which maintain the CENH3 mark. During cell-cycle dependent loading of CENH3, the CENH3 depleted chromosomes of the HI compete poorly with wild-type ones. Targeted depletion of CenH3^Cid^ in Drosophila produced haploids (Raychaudhuri et al., 2012). Furthermore, partially depleted centromeres could persist for many generations. We hypothesize that mass action kinetics favors centromeres with high density of CENH3 nucleosomes (Bodor et al., 2014). Notably, the depleted chromosomes maintain some identity, probably because removal of CENH3 or its associated factors is incomplete. When both parents provide CENH3-depleted centromeres (self cross or HIxHI cross), the absence of competition between centromeres enables the CENH3 chromatin to regain a functional homeostatic level during embryogenesis. In the WT, CENH3 density-dependent competition may help maintain dominance of the centromere over potential ectopic loci seeded by CENH3.

The role of CENH3 density in GE is demonstrated by the behavior of *cenh3-1* gametes produced by the *CENH3/cenh3-1* heterozygote. The chromosomes transmitted by these gametes display compromised kinetochore assembly, as in the CENH3-ts HI, and cause HI. Efficient GE by this mechanism was recently reported in maize (Wang et al., 2021). In addition, it may explain why a wheat HI CENH3 mutant, a restored-frameshift allele, resulted in GE when the heterozygote, but not the homozygote is selfed (Lv et al., 2020). In conclusion, rapid depletion of CENH3 can result in centromeric failure as demonstrated in fly and human (Hoffmann et al., 2016; Ly et al., 2017; Raychaudhuri et al., 2012).

CenH3-depleted centromeres can recover. In a GE cross, CENH3 depleted chromosomes missegregate frequently, leading to uniparental GE in a fraction of the progeny. Many zygotes eventually form diploids or aneuploids demonstrating resilience of the depleted centromeres. This resilience is accompanied by gradual acquisition of critical CENH3 density, providing a model for centromere recovery. CENP-A depleted centromeres in HeLa cells were competent and partly reassembled CENP-A (Mitra et al., 2020), whereas gametic depletion in flies results in loss of centromere function in embryos (Raychaudhuri et al., 2012). GE in HI crosses appears stochastic: it varies between and within embryos. Embryos chimeric for the expression of a maternal (HI) marker are evident at all developmental stages, suggesting that shoot apical meristems in haploid, diploid, and aneuploid sectors can give rise to corresponding plant types (Tan et al., 2015). Because different CENH3 variants form an allelic series varying in GE efficiency (Kuppu et al., 2020; Maheshwari et al., 2015), the epigenetic strength of centromeric identity, and the potential for recovery must vary proportional with the removal efficiency of each variant. At the same time, our observations and the high viability and absence of GE in HI x HI crosses indicate that when both parents contribute centromeres depleted below a critical level, CENH3 is reloaded in a manner sufficient for function. In conclusion, HI centromeres retain a weak, but distinct epigenetic memory.

An epigenetic factor regulates CENH3. A genetic screen identified natural variation in GE efficiency of the *GFP-ts* strain. Loss of epigenetic factor VIM1, which can ubiquitinate CENH3 in vitro and affects both DNA methylation and chromocenter size (Kraft et al., 2008; Woo et al., 2007), dramatically increased GE efficiency. Loss of VIM1 had an effect when either parent was the *vim1* mutant, suggesting postzygotic activity. This could be explained if a critical level of VIM1 is needed to stabilize CENH3 or facilitate loading, either through a stabilizing ubiquitin mark, or through differential DNA methylation. Post-translational modifications, such as sumoylation and phosphorylation in plants (Feng et al., 2020; Mérai et al., 2014; Zhang et al., 2005), methylation and ubiquitination in yeast (Ranjitkar et al., 2010; Samel et al., 2012), are often important or essential (Niikura et al., 2015, 2019).

Wang and Dawe (2018) proposed that competition between centromeres of different sizes results in GE. Most somatic cells of the HI display centromeres of comparable size (Kuppu et al., 2020) and specificity to the WT (Maheshwari et al., 2017). Furthermore, centromeres that appear much smaller than those of the WT, such as those observed when CENH3^G83E^ is used as male in a cross to the WT (Figure S5C), do not lead to measurable GE (Kuppu et al., 2020). It is unclear, however, whether a system based on dynamic CENH3 depletion constitutes a proper test for this hypothesis.

The property of CENH3 mutations described here could have interesting evolutionary implications. There is good evidence for the requirement of an optimal, species-specific CENH3 structure (Henikoff et al., 2001). In arabidopsis, evolutionary divergence of the complementing CENH3 results in increasing GE efficiency (Maheshwari et al., 2015), suggesting a progressively more severe mismatch with co-adapted factors. Changes in CENH3 structure may expose CENH3 to a surveillance system whose presence is well established in yeast and humans (Hewawasam et al., 2010; Niikura et al., 2019; Ohkuni et al., 2016; Ranjitkar et al., 2010). In this context, species differences are likely. The efficient centromeric function in crosses between CENH3-dependent HIs, suggests that evolution of species with sub-efficient CENH3 function is possible. For example, the CENH3^G83E^ mutation resulting in GE, could persist at low frequencies because it is recessive. Rarely, it may become fixed in a geographically isolated sub-population without affecting short-term fitness, as suggested by surveys (Kuppu et al., 2015). Lethality in the HI x WT cross should reinforce speciation by establishing a postzygotic barrier. The presence of this CENH3 variant, however, could expose these individuals to increased threat by neocentromeres if the reduced difference between the centromere and secondary CENH3 loci lessens the mass-action differential effect (Fig.9). Selection against the resulting genome instability would favor compensatory changes in CENH3 and interacting kinetochore proteins, and perhaps help explain the rapid evolution of CENH3 (Henikoff et al., 2001).

**Figure 9.**
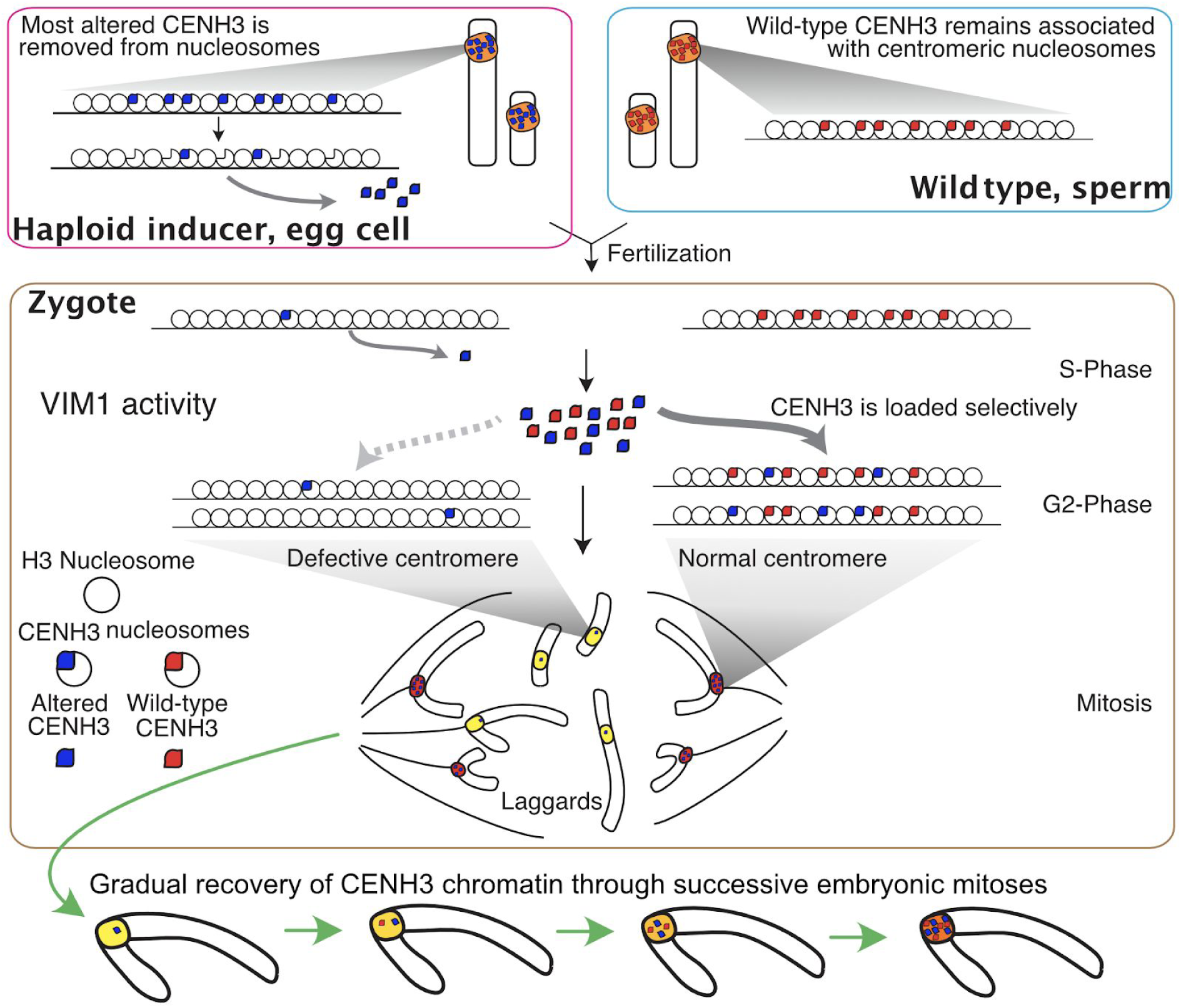
Model for haploid induction. Altered CENH3 is removed selectively from chromatin at the mature-egg stage. Wild-type CENH3 is not. In the cell divisions following hybridization, CENH3-depleted “weak” centromeres must compete for CENH3 loading with wild-type centromeres. Due to mass action kinetics (Bodor et al., 2014), wild-type centromeres are favored and load CENH3 preferentially. In the ensuing mitosis, the HI chromosomes missegregate due to their “weak” centromeres. The recessive action of altered CENH3 is accounted for by the persistence of wild-type CENH3. We further hypothesize that selective removal results from a genomic surveillance mechanism that eliminates defective or misplaced CENH3 molecules.

The difficulty in replicating the arabidopsis HI system in crops (Lv et al., 2020; Wang et al., 2021), could also be explained by species-specific variation in quantitative or developmental features of mechanisms that regulate CENH3 deposition and stability. The GFP-ts alteration is effective in arabidopsis, but probably lethal in maize (Wang et al., 2021). At the same time, transmission of a null *cenh3* allele yields 5% haploids in maize, less than 1% in arabidopsis, and none in wheat (Lv et al., 2020; Wang et al., 2021). Designing an efficient HI in each species may require different CENH3 modifications, as different constraints could apply to the function of CENH3 variants. Direct manipulation of CENH3 removal in the gametes may constitute a more general strategy.

In conclusion, our analysis of CENH3-mediated haploid induction in *A. thaliana* demonstrates its dependency on selective destabilization of HI-CENH3 variants during fertilization. The resulting differences in centromere loading efficiency provide insight into mechanisms that maintain the epigenetic memory of centromeric chromatin. Together, these findings provide a firm basis for further mechanistic insights and a framework for evaluating mechanisms of GE in biotechnology and during interspecific hybridization.

## Materials and Methods

### Plant growth and materials

Arabidopsis plants were grown under a 16 hr light/8 hr dark conditions 20°C in the Controlled Environment Facility at UC Davis. Unless mentioned otherwise, all lines (except lines carrying null alleles of *CenH3*) used in the study were homozygous for the genotype of interest. The lines carrying *cenh3-1* and *cenh3-3* null allele were used in heterozygous state with wild-type *CENH3* allele. The following lines have been previously described: *Cenh3/cenh3-1*, *GFP-ts*, *GFP-ts;CENH3, GFP-CENH3*, *tailswap-CENH3* (Ravi and Chan, 2010; Ravi et al., 2010), Simon Lab); *Lo*CENH3 and *Zm*CENH3 (Maheshwari et al., 2015, 2017), Comai lab), CenH3^G83E^(M4) and *CenH3^G173E^(M47)*, *cenh3-2^A86V (^Kuppu et al., 2015^)^*, Britt lab), *vim-1-2* (SALK 000930c) and Bor-4 (CS22591) (Woo et al., 2007) from Arabidopsis Biological Resource Center. Lines carrying *proEC1-H2B-RFP* and *proHTR10-HTR10-RFP (Ingouff et al., 2010)* was kindly provided by Berger lab (GMI, Austria); The line carrying *pRPS5A-H2B-tdTomato* (Maruyama et al., 2013) was kindly provided by Tetsuya Higashiyama (Nagoya University); A population of *4x GFP-ts* segregating for *cenh3-1* and *CENH3* alleles were kindly provided by Nico De Storme, Geelen lab (University of Ghent). Col-0 (2x), L*er gl-1*(2x) (Simon Chan lab Stocks) and L*er* 4x (Comai lab) were used as wild-type strains in the context of *CENH3*. *GFP-CENP-C, Nuf2-GFP* and *NDC80-GFP* (unpublished marker lines from Simon lab). The following lines employed in this study were generated by the floral dip method for Arabidopsis using corresponding constructs generated with p*CAMBIA1300* or *pCAMBIA3300* vectors in the late Dr. Simon Chan lab and the Comai lab: *proFWA-H2B-eCFP*, *proEC1-H2B-CFP*, *proCENP-CTagRFP-CENP-C* (mentioned as *RFP-CENP-C* in the text), *proNuf2-Nuf2-TagRFP* (mentioned as *Nuf2-RFP* in the text) and *proAct2-NLS-GFP-GUS*. A new null of *cenh3-3* (23bp deletion from +1177 bp to +1199bp) generated in L*er*(*gl1)* ecotype using *pKAMA-ITACHI Red CRISPR-Cas9* system (Tsutsui and Higashiyama, 2017) with single guide (5’ CCCCTCCCCAAATCAATCGT 3’) targeting 8th exon. The transgenic locus (Cas-9 and guide) were segregated out in the T2 generation and T3 generation lines carrying *cenh3-3* allele was used for the experiments. Additional details of all plasmid constructs and strains used in this study will be provided upon request.

### Emasculation and pollination

Mature buds were identified and emasculated a day before pollen shedding. On the following day, either the ovules from the emasculated buds were directly imaged or pollinated with appropriate male genotypes for imaging double-fertilization events, various stages of embryo development or for collecting seed for further progeny analysis.

#### Staging flower buds for imaging

For recording the chronological dynamics of GFP-ts during female gamete development on a single inflorescence axis, unopened buds were given negative(- sign) numbers and open flowers were given positive(+ sign) numbers (Figure 2). In our growth conditions, each healthy inflorescence branch of arabidopsis in early to mid-stages of development produces more than one flower per day. Hence, we used numerical nomenclature with minus and plus signs (Borg et al., 2020) to specify relative stages of the buds and flowers instead of standard staging nomenclature for Arabidopsis based on overall flower (Smyth et al., 1990) or ovule development (Schneitz et al., 1995). On a given inflorescence axis, the -1 stage being the matured bud and -2 stage is chronologically immediate but younger. In contrast, the +1 stage is chronologically young but the open flower and +2 stage is chronologically older and open flower (Fig. 2A). The buds in the -2 stage carry a mixture of undifferentiated gametes and differentiated egg cells. whereas -1 stage buds predominantly differentiated egg cells. The +1 and +2 stage flowers carry ovules with differentiated egg and central cells which are ready for fertilization.

### Imaging early stages of of zygote, embryo and endosperm development

For post fertilization analysis, the egg-cell chromatin was labeled with Histone H2B-CFP (At5g22880) or H2B-RFP (Ingouff et al., 2010) and the central cell or endosperm is marked by H2B-CFP (At5g22880) fusion in independent lines. These fusion proteins are driven by *EC1* (a egg-cell specific promoter, (Ingouff et al., 2007) and *FWA* (a central-cell and endosperm specific promoter, (Kinoshita et al., 2004) promoters. (Figure 1C). The sperm chromatin was marked by H2B-tdTomato or HTR10-RFP (Histone H3 variant) fusions driven respectively by *RPS5A* promoter (Maruyama et al., 2013) or sperm cell-specific *HTR10* promoter (Ingouff et al., 2010) (Figure 1C). Following fertilization, in *pEC1-H2B-CFP* X *pHTR10-HTR10-RFP* crosses, the zygotes were identified by colocalization of H2B-CFP and HTR10-RFP signals whereas endosperm were exclusively marked by HTR10-RFP from the male. Similarly in *pFWA-H2B-CFP* X *pHTR10-HTR10-RFP* crosses, endosperm was identified by colocalization of H2B-CFP and HTR10-RFP signals whereas zygotes were marked exclusively by HTR10-RFP. Similar system was used to identify the endosperm and embryo while employing *pRPS5A-H2B-tdTomato* expressing males (Figure 1C).

### Ovule and embryo dissection

At selected time points, ovules from individual pistils were dissected out using insulin needles directly into a drop of mounting media (1X PBS and 50% glycerol) on the glass slides and coverslip was gently placed on top of it and corners were sealed with nail polish. The volume of mounting media was found to be critical for imaging. We typically use 15-25μl volume/22×22mm coverglass and the volume depends on the quantity and size(age) of the ovule. More volume increases the thickness of the imaging plane in the z-axis which results in poor signal quality. In contrast, below a threshold volume, the ovules may get smashed and gametic or endosperm nuclei may be disfigured or released through the micropyle. Embryos were manually dissected from the fertilized ovules from 2 to 14-DAP using a fine tungsten needle while immersed in 0.1X PBS solution under a stereo microscope. While handling 4-cell stage embryos (two cells in the embryo proper, the future embryo, and two cells in suspensor), often the bottom cell or both cells in the suspensor were severed during dissection. The dissected embryos were transferred to glass slides with ∼5μl of the same buffer using a fine glass tube or 10μl plastic pipette tip coated with BSA (100mg/ml). Leaving 2-3μl of buffer with embryos, the rest of the medium was quickly removed and the mounting media (3-5μl of 1XPBS, 50% glycerol with 1μg/ml DAPI) was immediately added. A coverslip was gently placed on the top of the samples for direct observation. Mounted embryo or ovule samples were imaged on the same day of preparation. For immunostaining, dissected embryos were transferred to a glass slide in a drop of 0.1x PBS and proceeded further as described below.

### Immunostaining of nucleus and embryos

Nuclei were extracted in PBS by chopping the formaldehyde-fixed 2-3 young flower buds with a sharp razor blade. Chopped tissue mass was resuspended in 1ml cold PBS and filtered through a 40μm strainer. Nuclei in the filtrate was concentrated by centrifugation (250 RCF for 5 minutes). Leaving ∼15 μl of the supernatant along with the nuclei pellet, the rest of the supernatant was gently removed and discarded. 1-2μl of nuclei suspension was used for immunostaining. Immunostaining on isolated nucleus, dissected embryos and whole-mount ovules was performed according to (She and Baroux, 2014) with minor modifications. Immunostained samples were mounted with Prolong^TM^ Gold antifade with DAPI (Thermo Fisher) before imaging. Antibodies used in immunostaining: Primary: Rabbit CENH3 (Talbert et al., 2002) (1:2000), GFP#ab6556 (1:400), H3K4Me3 (1:500 #07-473-Milipore), H3K9me2 (1:200# ab1220), H3S10ph, 1:100 #ab14955). Secondary: Alexa fluor 405, 488, 594, 647 from Invitrogen used in (1:100 to 1:500 dilutions).

### GUS staining

GUS staining of embryo and seedling was carried out as described (Sundaresan et al., 1995). Dissected embryos (see above) were directly transferred to the GUS staining solution.

### Seed death and Haploid induction

In the GE crosses, seed death can be used as an indirect measure, which is proportional to haploid induction efficiency (Kuppu et al., 2020). For most of the experiments, crossed seeds collected on an individual silique (fruit) basis and the numbers were pooled following the quantification. For the seed death shown Figure 8D, for all of the cross combinations (except Wild-type(Col-0) x *GFP-ts* (n=87) and *GFP-CENH3* x *M47(CenH3^G173E^)*(n=48)) at least more than 240 and upto 923 seeds were accounted. The haploid frequency was scored by phenotyping the progeny as described in (Kuppu et al., 2015; Ravi and Chan, 2010).

### Imaging

Fluorescence images of nucleus, ovule, embryo and pollen samples were captured as 3-D objects using Applied Precision DeltaVision deconvolution or spinning disk confocal microscope in MCB light microscope imaging facility (UC Davis) and Zeiss LSM710 confocal microscope in department of Plant Biology (UC Davis). Images were captured at 60x magnification with Z-stacks (with a step size of 0.2μm for embryos and up to 1µm for ovules for initial screening and 0.2-0.5μm on selected images). One selected prometaphase sample (Figure 5C) was imaged with Nikon Structured Illumination Microscope for higher resolution. Zeiss Discovery v20 stereoscope was used to capture seeds, seedlings, GUS stained samples, and images processed with Zen lite software (Zeiss).

#### Image analysis

Images captured with Applied Precision DeltaVision deconvolution microscope were deconvolved with DeltaVision *softWoRx*™ before analysis. All 3-D images were analyzed with Imaris or SlideBook (only for 3i SDC image acquisition) softwares. If needed, series of consecutive z-planes were analyzed for resolving overlapping signals. On selected 3-D images, only z-stacks with cells or tissue of interest were selected and transformed into two-dimensional (2-D) using <u>m</u>aximum <u>i</u>ntensity <u>p</u>rojection (MIP) method. Selected images were processed using Adobe Photoshop and figures were assembled with Affinity Designer. While processing and analyzing images, brightness and contrast were altered and applied to the whole images in order to (1) reduce the general background (2) make the fainter signals visible relative to the brighter signals in the same image (3) to reduce the background autofluorescence in the ovule whole mounts.

To generate all figures in the manuscript, we selected representative images from each experiment, but whenever possible we chose images that had no overlapping centromere or kinetochore signals upon transforming into 2-D images (MIP). The same criteria were used to choose the images for signal intensity analysis presented in Figures 4, 6, 7, 8, S3 and S5. Centromeric signal (fluorescence) intensity of GFP-ts, GFP-CENH3 or CENH3 (by immunostaining) was measured using Softworx Explorer (Applied Precision Inc.). By scanning through the z-axis for each 3-D image, a 5×5 or 6×6 pixels area was selected with the region of interest aligned at the center and the intensity maximum from the point spread function for each signal was recorded <u>(Joglekar et al. 2008)</u>. Background noise was removed by selecting the same size region next to the ROI for each signal. Given that the analyzed cells may be in different stages of cell-cycle, collected centromeric fluorescence values (AU.) were normalized within every analyzed cell and expressed as relative fluorescent intensity. Qualitative patterns of centromeric GFP signal intensity were assigned by visual inspection of individual nuclei in using Imaris software. Graphs were generated with R-studio. The egg and central cells were readily identified by the expression of H2B-CFP marker driven by promoters EC1 and FWA respectively in addition to their position in the ovules guided by autofluorescence of ovule (Gooh et al., 2015; Ingouff et al., 2010). The centromeric GFP-ts signals in the undifferentiated gametes were recognized by presence of GFP signals in a group of 5 (= gametic chromosome number) in the mid-sections towards the micropylar end of the ovule which is otherwise free of somatic cells (Figure S2A). The CENH3 immuno signals in presumptive central cells in the wild-type background are recognized by its proximity to the egg egg cell and in the mid-sections of the ovule (Figure 2L).

## Author Contributions

M.P.A.M. planned and conducted research, interpreted results, wrote the manuscript, R.M. provided strains, and with R.B. participated in a part of the experimental design, and edited the manuscript, E-H.T. provided the seedling chimerism analysis, S.K. and A.B. provided genetic material, S.S.W.C. started this project and initiated research on it^$^, L.C. planned and analyzed research, wrote the manuscript.

## Acknowledgements

We would like to thank Steven Henikoff and Paul B. Talbert for sharing generous aliquots of the CENH3 antibody, Frederic Berger for sharing *pEC1-H2B RFP* and *pHTR10-HTR10-RFP* lines, Tetsuya Higashiyama for sharing *pRPS5A-H2B-tdTomato* line, Tomokazu Kawashima for suggestion on tissue-specific promoters, Savithramma Dinesh-Kumar for sharing plasmids with *RFP* and *CFP* genes, Célia Jäger-Baroux for suggestions and sharing unpublished immunostaining protocol embedded embryos, and Isabelle Henry and Andreas Houben for critical reading of the manuscript. We would like to thank the MCB Light Microscopy Imaging Facility at UC Davis for providing infrastructure for fluorescence imaging and analysis. Funding: Research in the laboratory of LC was was supported by a sub-award from the CSIRO for the grant “Capturing Heterosis for smallholders: OPP1076280” from the BMGF (USA), and from grant GBMF3068 from the Gordon and Betty Moore Foundation and from the Howard Hughes Medical Institute (HHMI). S.W.L.C. was a Howard Hughes Medical Institute–Gordon and Betty Moore Foundation Investigator. Research by SK and ABB was supported by Rijk Zwaan Zaadteelten Zaadhandel B.V., East-West Seed International, and Syngenta. Research in the laboratory of RM is supported by grant (STARS/APR2019/BS/818/FS dt: 31.12.2019) from Ministry of Education, Govt of India(GoI) and Focus basic research in the Agriculture Nutrition Biotechnology Theme project (MLP0120 Np.31-2(281)/208-19/budget dt. 14.09.2019) from CSIR, Ministry of Science and Technology, GoI. RB was supported by UGC JRF PhD fellowship awarded by University Grants Commission(UGC).

