## Supplemental Figures for "Biased removal and loading of centromeric histone H3 during reproduction underlies uniparental genome elimination"

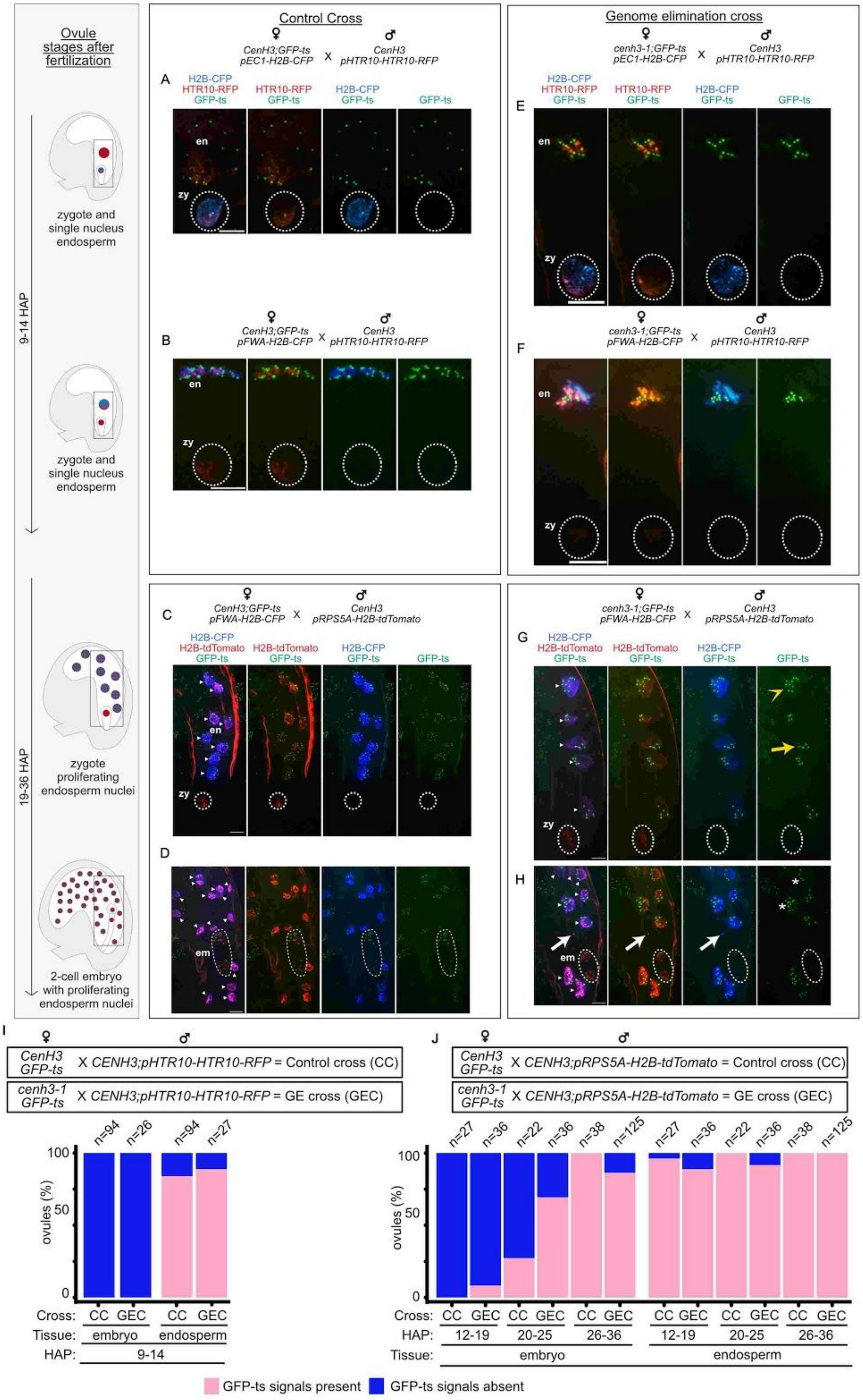
Ovule stages after fertilization
9-14 HAP
zygote and single nucleus endosperm
zygote and single nucleus endosperm
19-36 HAP
zygote proliferating endosperm nuclei
2-cell embryo with proliferating endosperm nuclei
Control Cross
CenH3;GFP-ts pEC1-H2B-CFP X CenH3 pHTR10-HTR10-RFP
A
H2B-CFP HTR10-RFP GFP-ts
HTR10-RFP GFP-ts
H2B-CFP GFP-ts
GFP-ts
en
zy
B
CenH3;GFP-ts pFWA-H2B-CFP X CenH3 pHTR10-HTR10-RFP
Genome elimination cross
cenh3-1;GFP-ts pEC1-H2B-CFP X CenH3 pHTR10-HTR10-RFP
E
F
cenh3-1;GFP-ts pFWA-H2B-CFP X CenH3 pHTR10-HTR10-RFP
C
CenH3;GFP-ts pFWA-H2B-CFP X CenH3 pRPS5A-H2B-tdTomato
H2B-CFP H2B-tdTomato GFP-ts
H2B-tdTomato GFP-ts
D
em
G
cenh3-1;GFP-ts pFWA-H2B-CFP X CenH3 pRPS5A-H2B-tdTomato
H
I
CenH3 GFP-ts X CENH3;pHTR10-HTR10-RFP = Control cross (CC)
cenh3-1 GFP-ts X CENH3;pHTR10-HTR10-RFP = GE cross (GEC)
n=94 n=26 n=94 n=27
ovules (%)
100
50
0
Cross: CC GEC CC GEC
Tissue: embryo endosperm
HAP: 9-14
J
CenH3 GFP-ts X CENH3;pRPS5A-H2B-tdTomato = Control cross (CC)
cenh3-1 GFP-ts X CENH3;pRPS5A-H2B-tdTomato = GE cross (GEC)
n=27 n=36 n=22 n=36 n=38 n=125 n=27 n=36 n=22 n=36 n=38 n=125
Cross: CC GEC CC GEC CC GEC CC GEC CC GEC CC GEC
HAP: 12-19 20-25 26-36 12-19 20-25 26-36
Tissue: embryo endosperm
GFP-ts signals present
GFP-ts signals absent

**Supplementary Figure 1. Haploid induction is marked by the presence of maternal GFP-ts on one of the parental chromatin sets ( x=5).**

Schematic representation depicting the regions of interest in the ovule stages shown in the control (A, B, C, D) and GE cross (E, F, G, H) images on the right. Lines expressing *pHTR10-RFP* (a sperm-specific marker) are used as male parent in A, B, E and F whereas lines expressing p*RPS5A-H2B-tdTomato,* a marker active in dividing cells, are used as male parent in C,D,G and H. Zygote (zy) and embryo (em) are marked with dotted white circles or ellipses, respectively. Endosperm is marked by "en" or white triangles. A yellow arrowhead and yellow arrow mark endosperm nuclei with, respectively, 10 and 5 centromeric GFP-ts signals in G. "*" marks the endosperm nuclei carrying bright and fainter centromeric GFP-ts signals in H. Quantification of presence and absence of GFP-ts signals in zygote and endosperm nuclei at early stages of development in control and genome elimination crosses (HI) with *pHTR10-HTR10-RFP* (I) or *pRPS5A-H2B-tdTomato* (J) as male parent. n=number of ovules. HAP: hours after pollination. Scale bars= 5µm.

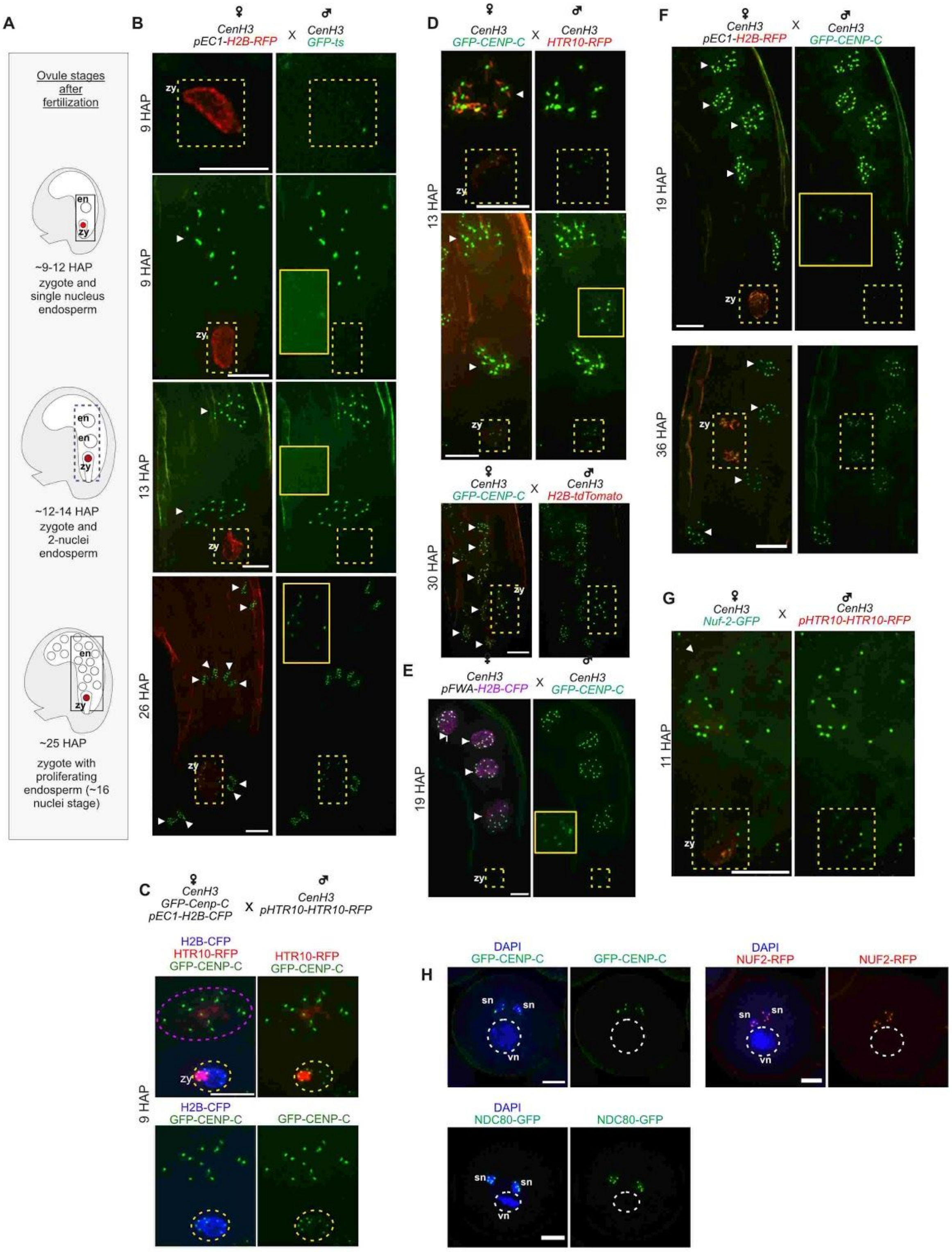
A
Ovule stages after fertilization
en
zy
~9-12 HAP zygote and single nucleus endosperm
~12-14 HAP zygote and 2-nuclei endosperm
~25 HAP zygote with proliferating endosperm (~16 nuclei stage)
B
♀ CenH3 pEC1-H2B-RFP
X
♂ CenH3 GFP-ts
9 HAP
13 HAP
26 HAP
C
♀ CenH3 GFP-Cenp-C pEC1-H2B-CFP
X
♂ CenH3 pHTR10-HTR10-RFP
H2B-CFP HTR10-RFP GFP-CENP-C
HTR10-RFP GFP-CENP-C
H2B-CFP GFP-CENP-C
GFP-CENP-C
D
♀ CenH3 GFP-CENP-C
X
♂ CenH3 HTR10-RFP
♀ CenH3 GFP-CENP-C
♂ CenH3 H2B-tdTomato
30 HAP
E
♀ CenH3 pFWA-H2B-CFP
♂ CenH3 GFP-CENP-C
19 HAP
F
♀ CenH3 pEC1-H2B-RFP
♂ CenH3 GFP-CENP-C
36 HAP
G
♀ CenH3 Nuf-2-GFP
♂ CenH3 pHTR10-HTR10-RFP
11 HAP
H
DAPI GFP-CENP-C
GFP-CENP-C
DAPI NUF2-RFP
NUF2-RFP
DAPI NDC80-GFP
NDC80-GFP
sn
vn

**Supplementary Figure 2. Loading dynamics of GFP-ts, and kinetochore proteins in embryo, endosperm and pollen.**
(A) Schematic representation depicting the regions of interest in the ovule stages shown in images B-G. Ovules regions in various stages of zygotic and endosperm development: (B) Dynamics of *CENH3;GFP-ts* as male parent (C) Loading of maternal GFP-CENP-C on paternal centromeres immediately after fertilization in both zygote (marked by maternal H2B-CFP and paternal HTR10-RFP) and endosperm (marked by paternal HTR10-RFP). Persistence of maternal GFP-CENP-C (D) and paternal GFP-CENP-C (E and F) in early stages of zygote and endosperm development. (G) Loading of maternal NUF-2-GFP on the paternal centromeres. The yellow dotted box or circle highlight the zygote. The same area is enlarged and enhanced in the inset with a solid yellow border in B, D, E and F. White solid triangle or dotted-purple circle (C) mark the endosperm nucleus. (H) Absence of kinetochore marker signals in the vegetative nuclei of mature pollen (white dotted circles). Individual pollen grains are demarcated by autofluorescence from the pollen wall. en: endosperm; zy: zygote; sn. Sperm nucleus; vn. vegetative nucleus. Scale bars=5μm.

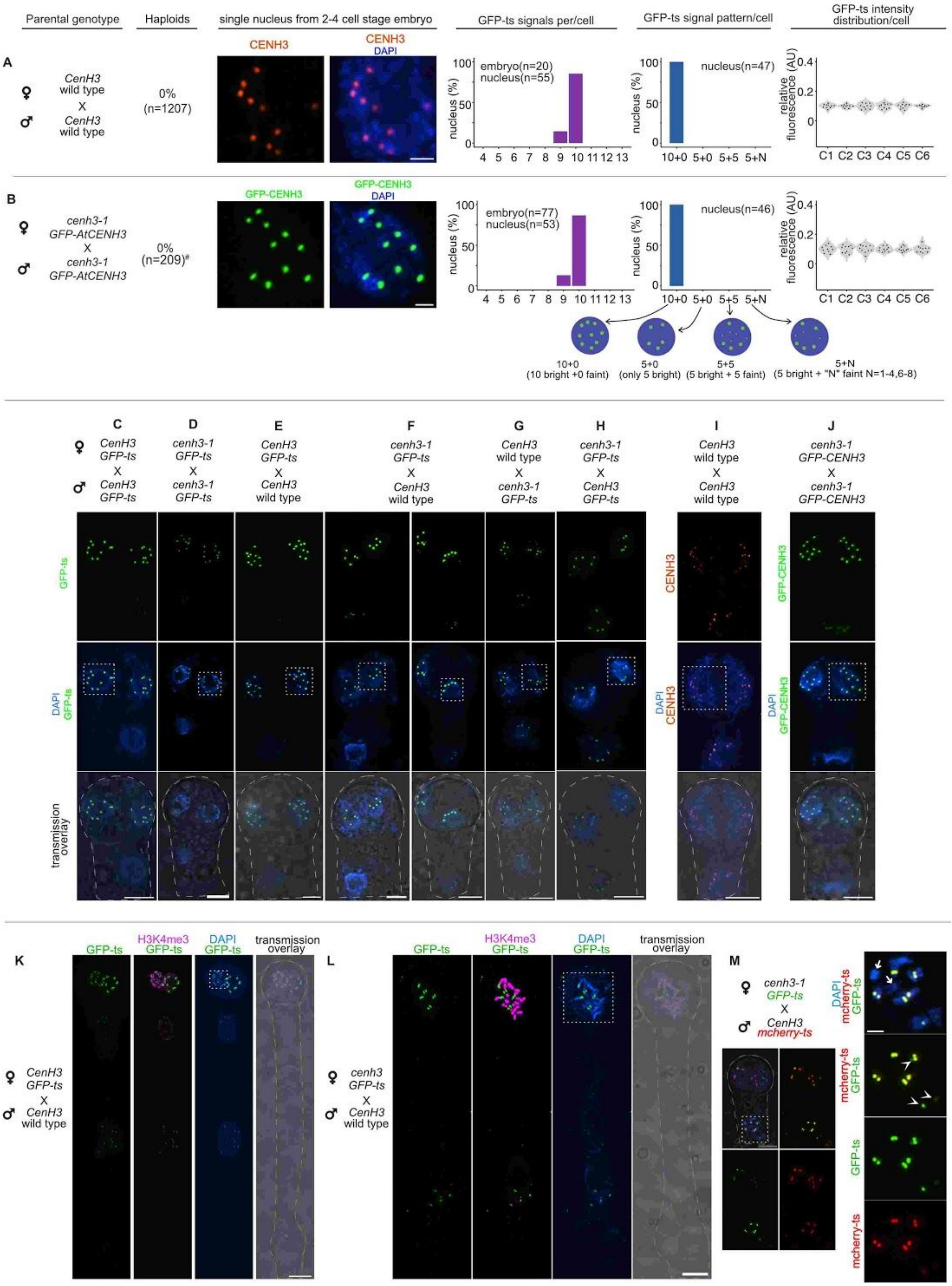

Parental genotype
Haploids
single nucleus from 2-4 cell stage embryo
GFP-ts signals per/cell
GFP-ts signal pattern/cell
GFP-ts intensity distribution/cell
A
CenH3 wild type
X
CenH3 wild type
0%
(n=1207)
CENH3
CENH3
DAPI
embryo(n=20)
nucleus(n=55)
nucleus(n=47)
nucleus (%)
relative fluorescence (AU)
B
cenh3-1 GFP-AtCENH3
X
cenh3-1 GFP-AtCENH3
0%
(n=209)#
GFP-CENH3
GFP-CENH3
DAPI
embryo(n=77)
nucleus(n=53)
nucleus(n=46)
10+0 5+0 5+5 5+N
C1 C2 C3 C4 C5 C6
10+0
(10 bright +0 faint)
5+0
(only 5 bright)
5+5
(5 bright + 5 faint)
5+N
(5 bright + "N" faint N=1-4,6-8)
C
CenH3 GFP-ts
X
CenH3 GFP-ts
D
cenh3-1 GFP-ts
X
cenh3-1 GFP-ts
E
CenH3 GFP-ts
X
CenH3 wild type
F
cenh3-1 GFP-ts
X
CenH3 wild type
G
CenH3 wild type
X
cenh3-1 GFP-ts
H
cenh3-1 GFP-ts
X
CenH3 GFP-ts
I
CenH3 wild type
X
CenH3 wild type
J
cenh3-1 GFP-CENH3
X
cenh3-1 GFP-CENH3
GFP-ts
DAPI GFP-ts
transmission overlay
CENH3
DAPI CENH3
GFP-CENH3
DAPI GFP-CENH3
K
CenH3 GFP-ts
X
CenH3 wild type
GFP-ts
H3K4me3 GFP-ts
DAPI GFP-ts
transmission overlay
L
cenh3 GFP-ts
X
CenH3 wild type
M
cenh3-1 GFP-ts
X
CenH3 mcherry-ts
DAPI mcherry-ts GFP-ts
mcherry-ts GFP-ts
GFP-ts
mcherry-ts

**Supplementary Figure 3. Biased loading of GFP-ts onto one set(x=5) of the parental centromeres at 2-4 cell stage hybrid embryos highlights genome elimination.**

Embryo nuclei from progeny of selfed (A) wild-type (CENH3 immunolocalization in red) and selfed (B) *cenh3-1* mutant complemented with the GFP-tagged-CENH3 construct (GFP-CENH3). The bar graph data displaying centromere signal patterns were obtained from a subset (carrying a euploid number of brighter GFP-ts signals) of total nuclei analyzed for each cross. Each column in the violin plot indicates relative GFP-ts signal intensity from six (C1-C6) nuclei. (C-L) Representative embryos after the second zygotic mitosis stage in both control and genome-elimination crosses. This material was used for the data shown in Figure 4 (A to I) and supplementary figure 4 (A and B) (selected single nuclei are highlighted with white dotted box). (M) Paternal *mcherry-ts* reflects the maternal GFP-ts loading pattern in a HI cross. White dotted area is magnified on the right highlighting the pre-mitotic stage. White arrowheads mark faint signals; white arrows mark condensed chromosomes without any centromeric signals. “#” Data from Ravi and Chan 2010. Scale bar= 1 µm(for single nucleus); 5 µm (for whole embryos).

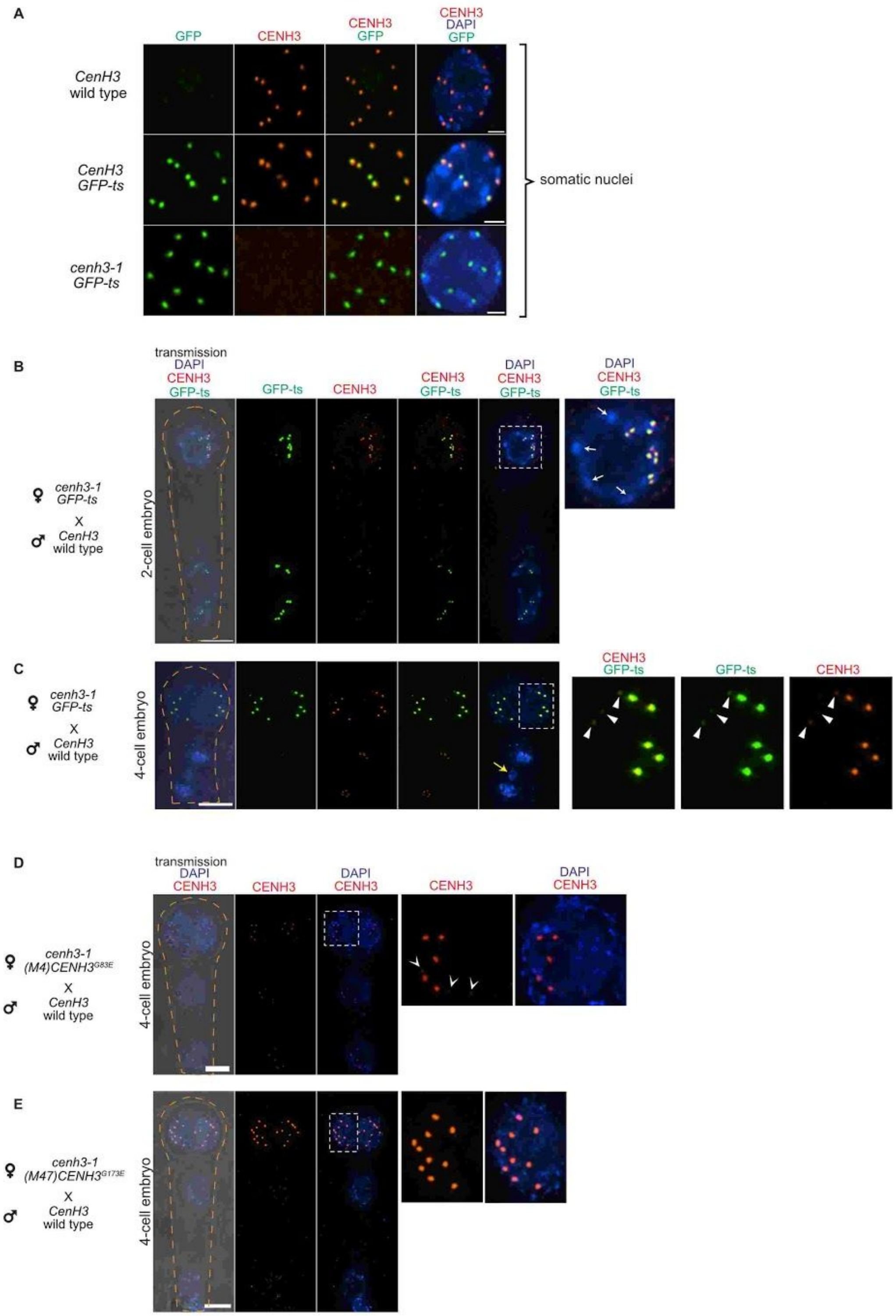
A
GFP
CENH3
CENH3
GFP
CENH3
DAPI
GFP
CenH3
wild type
CenH3
GFP-ts
cenh3-1
GFP-ts
somatic nuclei
B
transmission
DAPI
CENH3
GFP-ts
GFP-ts
CENH3
cenh3-1
GFP-ts
X
CenH3
wild type
2-cell embryo
C
cenh3-1
GFP-ts
X
CenH3
wild type
4-cell embryo
D
cenh3-1
(M4)CENH3G83E
X
CenH3
wild type
4-cell embryo
DAPI
CENH3
E
cenh3-1
(M47)CENH3G173E
X
CenH3
wild type
4-cell embryo

**Supplementary Figure 4: Native CENH3 and GFP tailswap-CENH3 load onto the same functional centromeres during haploid induction.**

(A) Representative nuclear staining with anti-CENH3 antibody demonstrates the absence of cross-reactivity with GFP-ts variant. Colocalization of GFP-ts and CENH3 on one parental set of centromeres in G2 nucleus of 2-cell (B) and 4-cell (C) stage embryo from GE cross. White arrows mark chromocenters without CENH3 or GFP-ts signals. White triangle marks colocalization of faint CENH3 signals with the faint GFP-ts signals. Yellow arrow marks laggard chromosomes without CENH3 or GFP-ts signals. Localization of CENH3 on one (D) or both parental sets of centromeres (E) respectively in GE cross (*CenH3*$^{G83E}$ x Wild-type) and control cross (*CenH3*$^{G173E}$ x Wild-type). White arrowhead (in D) marks fainter CENH3 signals. Scale bar= 1µm for the images in A and 5µm for the rest of the figure.

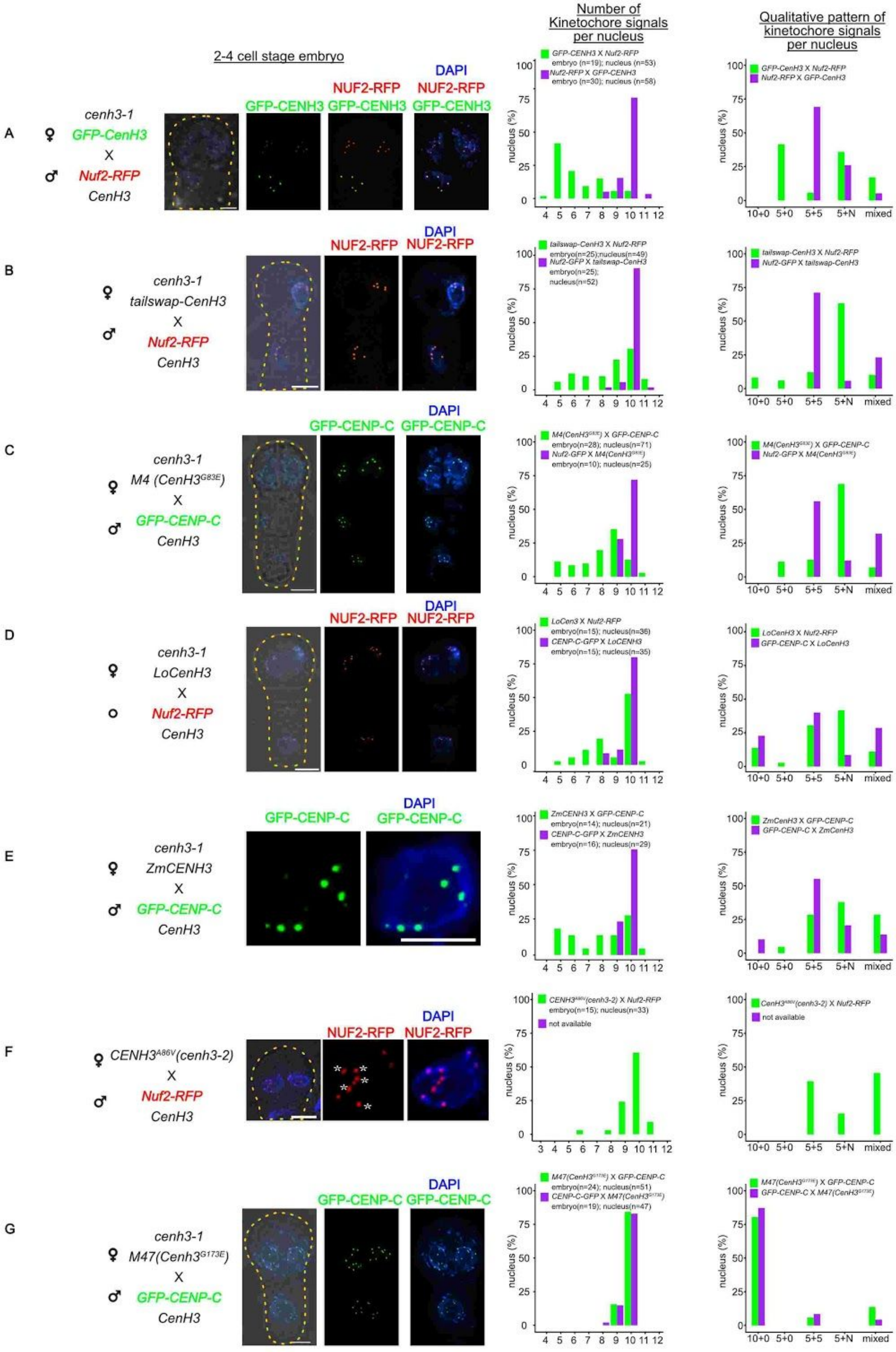
Number of Kinetochore signals per nucleus
Qualitative pattern of kinetochore signals per nucleus
2-4 cell stage embryo
A
cenh3-1
GFP-CenH3
X
Nuf2-RFP
CenH3
GFP-CENH3
NUF2-RFP GFP-CENH3
DAPI NUF2-RFP GFP-CENH3
B
cenh3-1
tailswap-CenH3
X
Nuf2-RFP
CenH3
NUF2-RFP
DAPI NUF2-RFP
C
cenh3-1
M4 (CenH3G83E)
X
GFP-CENP-C
CenH3
GFP-CENP-C
DAPI GFP-CENP-C
D
cenh3-1
LoCenH3
X
Nuf2-RFP
CenH3
NUF2-RFP
DAPI NUF2-RFP
E
cenh3-1
ZmCENH3
X
GFP-CENP-C
CenH3
GFP-CENP-C
DAPI GFP-CENP-C
F
CENH3A86V(cenh3-2)
X
Nuf2-RFP
CenH3
NUF2-RFP
DAPI NUF2-RFP
G
cenh3-1
M47(Cenh3G173E)
X
GFP-CENP-C
CenH3
GFP-CENP-C
DAPI GFP-CENP-C
nucleus (%)
10+0 5+0 5+5 5+N mixed
not available

**Supplementary Figure 5. Two to four cell stage embryos from the genome elimination cross involving different haploid inducer lines predominantly assemble five stronger kinetochores.**

Two to four cell stage embryos from the genome-elimination crosses (A-F) or control cross (G) involving multiple CENH3 variants. The images of embryos and nuclei are shown only for the crosses where the CENH3 variant was used as female. GFP-CENP-C or NUF2-RFP expressing lines in the *CENH3 +/+* background used as a male line. Bar graphs in the right indicate the counts of kinetochore signals per nuclei of the respective genotypes shown on the left. The number of embryos and nuclei observed for each cross and used to generate bar graphs is shown in parenthesis. The “x+x” pattern indicates the number of bright + faint signal patterns. For eg. in the pattern 5+N, N=1,2,3,4,6. Mixed pattern indicates nucleus with a variable number of bright signals + faint signals that do not follow the expected euploid pattern(10+0, 5+0, 5+5, 5+N). Scale bar=5µm.

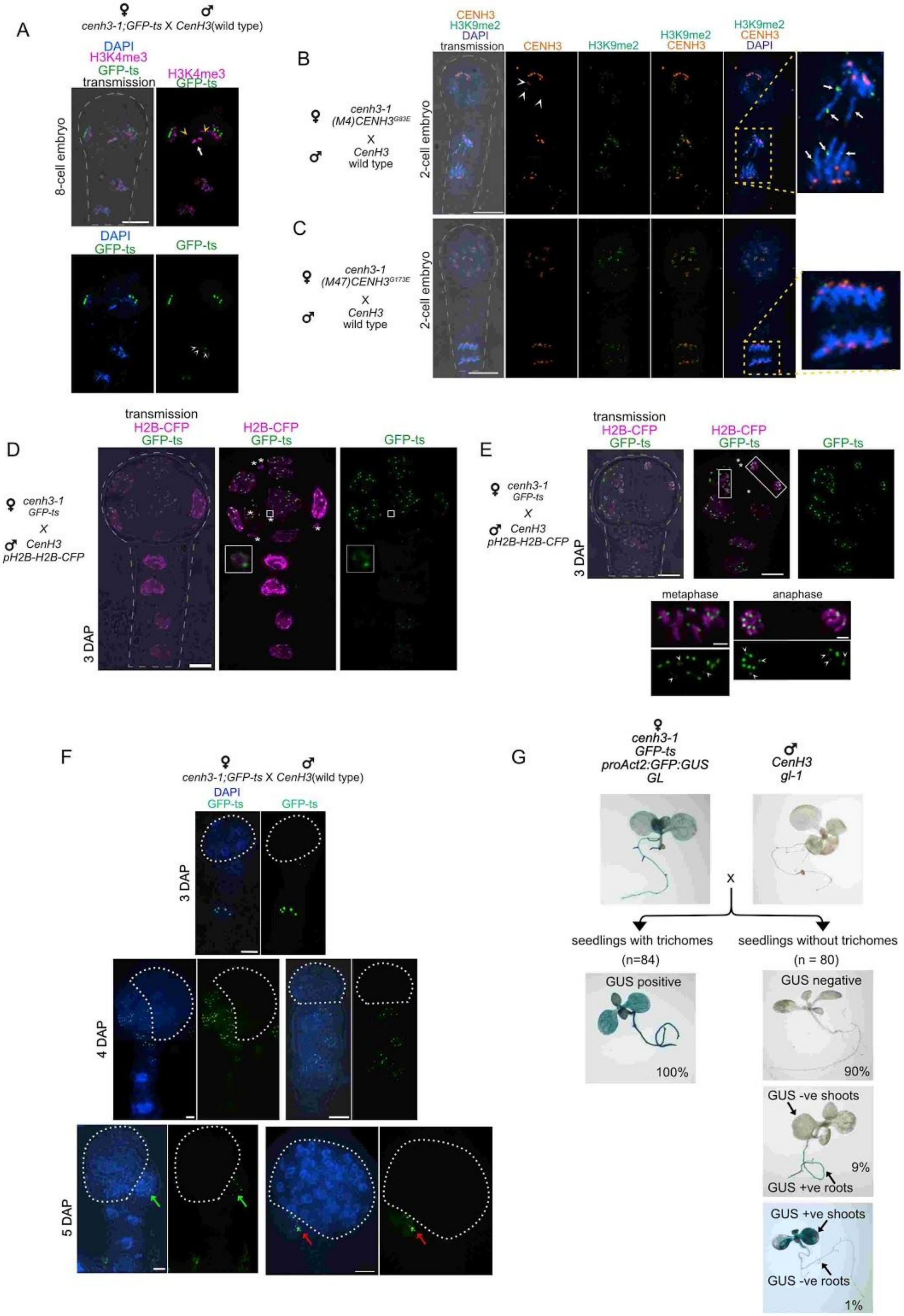
A
♀ ♂
cenh3-1;GFP-ts X CenH3(wild type)
DAPI
H3K4me3
GFP-ts
transmission
H3K4me3
GFP-ts
8-cell embryo
DAPI
GFP-ts
GFP-ts
B
CENH3
H3K9me2
DAPI
transmission
CENH3
H3K9me2
H3K9me2
CENH3
H3K9me2
CENH3
DAPI
♀ cenh3-1
(M4)CENH3$^{G83E}$
X
♂ CenH3
wild type
2-cell embryo
C
♀ cenh3-1
(M47)CENH3$^{G173E}$
X
♂ CenH3
wild type
2-cell embryo
D
♀ cenh3-1
GFP-ts
X
♂ CenH3
pH2B-H2B-CFP
transmission
H2B-CFP
GFP-ts
H2B-CFP
GFP-ts
GFP-ts
3 DAP
E
♀ cenh3-1
GFP-ts
X
♂ CenH3
pH2B-H2B-CFP
3 DAP
transmission
H2B-CFP
GFP-ts
H2B-CFP
GFP-ts
GFP-ts
metaphase
anaphase
F
♀ ♂
cenh3-1;GFP-ts X CenH3(wild type)
DAPI
GFP-ts
GFP-ts
3 DAP
4 DAP
5 DAP
G
♀
cenh3-1
GFP-ts
proAct2:GFP:GUS
GL
♂
CenH3
gl-1
X
seedlings with trichomes
(n=84)
seedlings without trichomes
(n = 80)
GUS positive
100%
GUS negative
90%
GUS -ve shoots
GUS +ve roots
9%
GUS +ve shoots
GUS -ve roots
1%

**Supplementary Figure 6. Fate of HI chromosomes in developing embryos.**

Early embryonic mitosis in GE (A, B, D and E) and control crosses (C). (A) H3H4me3 staining highlights the unstable HI parent chromosomes from an 8-cell stage embryo (only partial sections are shown). (B) Normal localization of H3K9me2 and biased (uniparental) localization of CENH3 on the centromeres of embryonic cells at G2 (top, embryo proper) and anaphase (bottom, suspensor cell). (C) Normal localization of H3K9me2 and CENH3 on the centromeres of embryonic cells at G2 (top, embryo proper) and anaphase (bottom, suspensor cell). (D) Embryo from an elimination cross from ovules 3 DAP showing the presence of multiple micronuclei. Inset: micronuclei with faint GFP-ts signal. (E) Embryo containing multiple micronuclei with metaphase and anaphase stage cells highlighted (rectangle). Note that both metaphase and anaphase cells display bright and faint GFP-ts centromeric signals. The chromosomes with fainter signals appear to segregate normally in the latter along with chromosomes with brighter signals. (A, B, D and E) Yellow arrowhead: chromatin bridge; white arrow: laggard chromosomes; white arrowheads: faint GFPts or CENH3 signals; "*" marks micronuclei. (F) Chimeric distribution of GFP-ts signals in two to five day old embryos. Cells lacking signals or displaying fainter signals in the embryo proper are highlighted (dotted white lines). Green arrow: A GFP-ts positive cell in an otherwise GFP-ts negative abnormal embryo, suggestive of aneuploidy. Red arrow: A GFP-ts positive sector in an otherwise GFP-ts negative embryo displaying normal development, suggestive of haploidy. (G) GUS staining revealing the chimeric nature genome elimination at a whole seedling level. Seedlings with trichomes represent hybrid diploid/ aneuploid progeny and without trichomes are predominantly haploid. n= number of seedlings. Scale bar= 5 μm for A-F.
